# STX4 is indispensable for mitochondrial homeostasis in skeletal muscle

**DOI:** 10.1101/2025.09.02.673840

**Authors:** Joseph M. Hoolachan, Rekha Balakrishnan, Erika M. McCown, Karla E. Merz, Chunxue Zhou, Elizabeth Bloom-Saldana, Patrick T. Fueger, Angelica Hamilton, Tali Kiperman, Ke Ma, Eunjin Oh, Lei Jiang, Patrick Pirrotte, Orian Shirihai, Debbie C. Thurmond

## Abstract

**Background:** Mitochondrial homeostasis is vital for optimal skeletal muscle integrity. Mitochondrial quality control (MQC) mechanisms that are essential for maintaining proper functions of mitochondria include mitochondrial biogenesis, dynamics and mitophagy. Previously, Syntaxin 4 (STX4) traditionally considered a cell surface protein known for glucose uptake in skeletal muscle, was also identified at the outer mitochondrial membrane. STX4 enrichment was sufficient to reverse Type 2 diabetes-associated mitochondrial damage in skeletal muscle by inactivation of mitochondrial fission. However, whether STX4 could modulate skeletal muscle mitochondrial homeostasis through MQC mechanisms involving mitochondrial biogenesis or mitophagy remains to be determined.

**Methods:** To determine the requirements of STX4 in mitochondrial structure, function and MQC processes of biogenesis and mitophagy, we implemented our in-house generated inducible skeletal muscle-specific STX4-knockout (skmSTX4-iKO) mice (*Stx4^fl/fl^; Tg(HSA-rtTA/TRE-Cre*)/B6) and STX4-depleted immortalized L6.GLUT4myc myotubes via siRNA knockdown (siSTX4).

**Results:** We found that non-obese skmSTX4-iKO male mice (>50% reduced STX4 abundance, Soleus and Gastrocnemius ***p<0.001, Tibialis anterior (TA) ****p<0.0001) developed insulin resistance (**p<0.01), together with reduced energy expenditure (AUC *p<0.05), respiratory exchange ratio (AUC **p<0.01), and grip strength (*p<0.05). STX4 ablation in muscle also impaired mitochondrial oxygen consumption rate (****p<0.0001). Mitochondrial morphological damage was heterogenous in STX4 depleted muscle, presenting with small fragmented mitochondria (****p<0.0001) and deceased electron transport chain (ETC) abundance (CI ***p<0.001, CII *p<0.05, CIV **p<0.01) in oxidative soleus muscle, while glycolytic TA fibers display enlarged swollen mitochondria (****p<0.0001) with no change in ETC abundance. Notably, >60% reduction of STX4 in siSTX4 L6.GLUT4myc myotubes (****p<0.0001) also decreased ETC abundance (CI ****p<0.0001, CII ****p<0.0001, CIV *p<0.05) without changes in mitochondrial glucose metabolism, as shown by [U-^13^C] glucose isotope tracing. For MQC, both skmSTX4-iKO male mice (*p<0.05) and siSTX4 L6.GLUT4myc myotubes (*p<0.05) showed decreased mitochondrial DNA levels alongside reduced mRNA expression of mitochondrial biogenesis genes *Ppargc1a* (PGC1-α, *p<0.05) and *Tfam* (*p<0.05) in skmSTX4-iKO soleus muscle and PGC1-α (mRNA *p<0.05, protein ***p<0.001), NRF1 (mRNA and protein *p<0.05) and *Tfam* (mRNA *p<0.05) in siSTX4 L6.GLUT4myc myotubes. Furthermore, live cell imaging using mt-Keima mitophagy biosensor in siSTX4 L6.GLUT4myc cells revealed significantly impaired mitochondrial turnover by mitophagy (*p<0.05) and mitochondria-lysosome colocalization (*p<0.05). STX4 depletion also reduced canonical mitophagy markers, PINK1 and PARKIN in both skmSTX4-iKO muscle (PARKIN *p<0.05, PINK1 **p<0.01) and siSTX4 L6.GLUT4myc myotubes (PARKIN ****p<0.0001, PINK1 *p<0.05).

**Conclusions:** Our study demonstrated STX4 as a key mitochondrial regulator required for mitochondrial homeostasis in skeletal muscle.

## 1 Introduction

Mitochondrial homeostasis to maintain a healthy functional mitochondrial pool is critical for skeletal muscle integrity. Mitochondria are dynamic organelles capable of altering their morphology, density and network distribution to adapt to metabolic demands and cellular processes [1]. For skeletal muscle, a tissue that accounts for 40% of total body mass [2] and >80% of whole-body glucose homeostasis [3], mitochondria play key roles in mediating myogenesis, muscle size, fiber heterogeneity, contraction and glucose and fatty acid metabolic processes [4].

Mitochondrial homeostasis requires a tightly coordinated set of mitochondrial quality control (MQC) mechanisms including dynamics, biogenesis and mitophagy [5–7]. Mitochondrial dynamics via fusion (elongated mitochondria) and fission (fragmented mitochondria) alters mitochondrial morphology to cope with metabolic demand [5,7,8], whilst the opposing processes of mitochondrial biogenesis (expansion of pre-existing mitochondrial population) and mitophagy (clearance of damaged and/or aging mitochondria) require a fine balance [5–7].

Mitochondrial biogenesis entails the coordination of transcription factors such as Nuclear respiratory factor 1 (NRF1) and Transcription factor A, Mitochondrial (TFAM) that promote mitochondrial protein expression for expansion of pre-existing mitochondria orchestrated by the master regulator Peroxisome proliferator-activated receptor-gamma coactivator (PGC1-α) [5,7,9]. In contrast, mitophagy, the clearance of damaged and/or aging mitochondria occurs via two autonomous processes:1) canonical PTEN-induced kinase 1 (PINK1)/PARKIN-mediated ubiquitination [7,10,11] and 2) non-canonical receptor-dependent mitochondrial membrane proteins in possession of microtubule-associated protein 1 light chain 1 β (LC3) interacting region (LIR) motifs [12]. LIR motifs directly connect the mitochondria with LC3-positive autophagosomes [10,12], such as FUNDC1 [13], AMBRA1 [14] and STX17 [12]. Recent findings indicate that PINK1/PARKIN-mediated mitophagy was observed predominantly in oxidative slow-twitch skeletal muscle types such as the soleus [11].

Impairments in MQC mechanisms across biogenesis [15], dynamics [8,16] and mitophagy [16] coincide with the presence of dysfunctional mitochondrial function during progression of insulin resistance from prediabetes towards Type 2 diabetes (T2D) [17] that culminates in damaged, vacuolized and fragmented mitochondria [8]. Previously, we made the novel discovery that Syntaxin 4 (STX4), traditionally considered a cell surface exocytosis protein in skeletal muscle critical for GLUT4-vesicle fusion in insulin-stimulated glucose uptake [18] and downregulated in T2D skeletal muscle, was additionally localized to the outer mitochondrial membrane (OMM) [8]. Furthermore, STX4 enrichment specifically in the skeletal muscle of high fat diet (HFD)-fed insulin resistant male mice reversed peripheral insulin resistance, increased energy expending efficiency in vivo, and improved the structure and function of mitochondria in skeletal muscle tissue and isolated myofibers ex vivo [8]. Of note, we identified STX4 function in modulating fission/fusion dynamics by promoting elongated mitochondrial networks via inactivation of fission protein Dynamin-related protein 1 (Drp1) with phosphorylation of S637 [8]. Given the beneficial effects of STX4 enrichment on HFD-associated mitochondrial damage, it remains to be determined whether STX4 is indispensable for mitochondrial structure/function and involved in MQC mechanisms (e.g., mitochondrial biogenesis and mitochondrial turnover by mitophagy).

The present study aimed to further explore STX4 function in skeletal muscle mitochondrial structure, activity and MQC using an inducible skeletal muscle-specific STX4 knockout (skmSTX4-iKO) mouse model together with a STX4-depleted muscle cell line. We hypothesized that STX4 was an important mitochondrial regulatory protein in skeletal muscle and modulates MQC.

## 2 Methods

### 2.1 Animal Studies

STX4 floxed mice (STX4^fl/fl^) were obtained from Dr. Sidney Whiteheart (University of Kentucky) [S1] and crossed with heterozygous human skeletal actin (HSA)-rtTA/TRE-Cre recombinase positive mice (Cre^+/-^) (JAX, #012433) to produce doxycycline (Dox)-inducible STX4^fl/fl^;Cre^+/-^ mice (C57BL6/J background). Male and female skmSTX4-iKO mice were generated by providing 2 mg/mL Dox in drinking water for 14 days for *Stx4* gene excision [8]. Non-Dox inducible control (CTRL) mice for skmSTX4-iKO experiments were on the STX4^fl/fl^;Cre^-/-^ background and provided 2 mg/mL Dox in the drinking water. All mice were housed on Pure-O’Cel bedding in groups of 3-5 and fed a standard maintenance chow diet (13% of kCal from fat; Picolab #5053, Fort Worth, TX) starting at 4 weeks-of-age to maintain a non-obese phenotype. All experimental procedures were approved by the Institutional Animal Care and Use Committee of City of Hope (Duarte, CA, USA; protocol #23041).

### 2.2 Immunoblot Analysis

Whole protein lysates of cell cultures and primary mouse tissue was lysed as previously described [8] Per standard procedures, proteins lysates were resolved using in-house generated 10-12% SDS-PAGE gels and transferred onto polyvinylidene difluoride (PVDF) membranes for immunoblotting. Membranes were incubated with primary antibodies (Table S1) overnight at 4°C. Secondary antibodies Goat Anti-Rabbit IgG (H L)-HRP Conjugate (Bio-Rad, #172-1019) or goat Anti-Mouse IgG (H L)-HRP Conjugate (Bio-Rad, #172-1011) were incubated for 1 hour at room temperature. Protein bands were detected using a Chemic-Doc Touch gel documentation system (Bio-Rad) with Amersham ECL (Cytiva, #RPN2106), ECL prime (Cytiva, #RPN2232) or Super Signal (Fisher Scientific, #34095). Protein levels were quantified using Image Lab 6.1 (Bio-Rad) and normalized against Tubulin housekeeping protein levels or ponceau staining for total protein lysate.

### 2.3 Insulin Tolerance Testing

20-week-old CTRL and skmSTX4-iKO mice underwent intraperitoneal insulin tolerance testing (IPITT) as previously described [8].

### 2.4 Glucose Tolerance Testing

20-week-old CTRL and skmSTX4-iKO mice underwent intraperitoneal glucose tolerance testing (IPGTT) as previously described [8].

### 2.5 Metabolic Caging and Body Composition Analysis

Following an initial overnight acclimatization, 26-week-old CTRL and skmSTX4-iKO male mice were metabolically phenotyped with data collected every hour starting from 6 am across 72 hours in an indirect calorimetry cage system (Promethion, Sable Systems, Las Vegas, NV, USA) that measured oxygen consumption (VO2), CO2 production (VCO2) to determine respiratory exchange rate (RER), energy expenditure, food and water consumption.

### 2.6 Oxygen Consumption Rate (OCR) Measurements

OCR measurements of flexor digitorium brevis (FDB) muscle harvested from both legs of 20-week-old CTRL and skmSTX4-iKO male mice were performed as previously described [8].

### 2.7 Grip Strength

Forelimb grip strength of the CTRL and skmSTX4-iKO male mice was performed using a grip strength meter (Harvard Apparatus) of three repeated measurements for each mouse tested as previously described [S2].

### 2.8 Transmission Electron Microscopy (TEM) and Mitochondrial Area Quantification

Preparation, fixation, embedment and imaging of soleus and TA muscle from skmSTX4-iKO mice using FEI Tecnai 12 transmission electron microscope equipped with a Gatan OneView CMOS camera was performed as previously described [8]. Mitochondrial area in four to six 6500x TEM images per mouse was calculated using Image J 1.54p software set at 600 distance in pixels:1 μm scale bar.

### 2.9 Citrate Synthase Assay

Citrate synthase activity was measured in 2 μg of gastrocnemius (GAS) homogenate as per manufactures instructions using the MitoCheck Citrate Synthase Activity Assay Kit (Cayman Chemicals, #701040) as previously described [S3].

### 2.10 Cell Culture

Rat immortalized L6.GLUT4myc skeletal muscle cells (Kerafast #ESK202-FP) myoblast growth and myotube differentiation were performed as previously described [8,9]. For insulin signaling, myotubes were serum- and glucose-starved in oxygenated FCB buffer (125 mM NaCl, 5 mM KCl, 1.8 mM CaCl2, 2.6 mM MgSO4, 25 mM HEPES, 2 mM pyruvate and 2% (wt/vol) BSA (Bovine Serum Albumin) (Fisher Scientific, #BP9704100) for 2 hours prior to 10 nM insulin (Sigma-Aldrich, #I5500) stimulation for 5 minutes.

### 2.11 Small interfering RNA (siRNA) Transfection

For STX4 knockdown studies, differentiation day (D)4 L6.GLUT4myc myotubes were transfected with 100 nM STX4-targeting siRNA (siSTX4) (ThermoFisher, Assay ID:s135769) sequence or non-specific targeting siRNA control (siCON) (ThermoFisher, #4390843) in a drop wise fashion using RNAiMax (ThermoFisher, #13778075) and Opti-MEM (Gibco, #31985088) overnight and fresh media added the next day as previously described [8]. These myotubes underwent 96 hours total siSTX4 knockdown to ensure optimal reduction of STX4 protein abundance.

### 2.12 Stable Isotope Tracing

L6.GLUT4myc myoblasts were cultured in 60 mm dishes and underwent myotube differentiation and 96 hours siCON or siSTX4 transfection at the D4 stage as mentioned above (see 2.11). For the final 6 hours the siCON and siSTX4 L6.GLUT4myc myotubes were incubated in 5 mM [U-^13^C]glucose (Sigma, #389374) labelled glucose-free DMEM (ThermoFisher, #11966025) alongside a non-radioactive labelled 5 mM glucose (Sigma, #G6152) DMEM control for analysis of glycolytic and tricarboxylic acid (TCA) cycle metabolites as previously described [8].

### 2.13 Mitochondrial Copy Number

Total DNA isolation from rat L6.GLUT4myc myotubes and mouse GAS muscle was performed per DNeasy Blood and Tissue Kit instructions (Qiagen, #69504). Mitochondrial DNA (mtDNA) copy number was calculated as the ratio of mtDNA represented by respective primers (Integrated DNA Technologies (IDT)) for mitochondrial genome encoded genes compared to nuclear genome encoded reference genes using the QuantiTect SYBR Green PCR Kit (Qiagen, #204143) for real-time qPCR. Rat and mouse primers are described in (Table S2) and were previously used [8,9].

### 2.14 RNA Isolation and qPCR

Total RNA isolation from L6.GLUT4myc cell line was performed per RNeasy Plus Mini Kit instructions (Qiagen, #74136) and from mouse primary tissue with TriReagent (Sigma, #93289) as previously described [8]. Complementary DNA (cDNA) was generated per High-Capacity cDNA Reverse Transcription Kit instructions (ThermoFisher, #4368814). The QuantiTect SYBR Green PCR Kit and primers (IDT, Table S2) were used for qPCR detection. Relative gene expression was quantified against the validated respective housekeeping gene controls for each model (IDT, Table S2).

### 2.15 Mito-Keima (mt-Keima) Visualization of Mitophagy

L6.GLUT4myc myoblasts at 80% confluence underwent a 6-hour transfection with mKeima-Red-Mito-7 (mt-Keima) plasmid DNA, a gift from Michael Davidson (Addgene, #56018) via drop wise fashion of L6 Avalanche (EZ Biosystems, #EZT-L600-1) and Opti-MEM according to manufacturer’s instructions. This was subsequently followed by a 42-hour transfection with siCON or siSTX4 via RNAiMax and Opti-MEM. The cells were then exposed to ± 10 μM Carbonyl cyanide 3-chlorophenylhydrazone (CCCP) (ThermoFisher, #L06932-MC), a known mitochondrial uncoupler [S4-5] for 4 hours to promote mitophagy. Cells were imaged using a Leica LSM 900 confocal fluorescence microscope (Zeiss) at 40x objective for quantification of mean fluorescence intensity using Zen Blue (Zeiss) on lysosomal-associated mitochondria (pH 4.5, ex: 561 nm) [S5], nuclei were stained by NucBlue (ThermoFisher #R37605).

### 2.16 MitoTracker-LysoTracker Visualization of Mitochondria-Lysosome Association

The siCON- or siSTX4-treated L6.GLUT4myc myotubes were exposed to ±10 μM CCCP for 4 hours and then treated with 100 nM MitoTracker Red (ThermoScientific, #M7510) and 50 nM LysoTracker Green (ThermoScientific, #L7526) for 30 minutes at 37°C in a 5% CO_2_ incubator. The myotubes were imaged using a LSM 900 fluorescence confocal microscope using a 63X oil-immersed objective; mitochondria (red)-lysosome (green) co-localization (yellow) was quantified using Pearson’s co-efficient in Zen Blue (Zeiss); nuclei were visualized with NucBlue.

### 2.17 Statistical Analyses

Statistical analyses were performed using GraphPad PRISM (San Diego, CA, version 10.1.0). Unpaired two-tailed Student’s *t*-test was used for single comparisons between genotype (CTRL vs skmSTX4-iKO) and cells (siCON vs siSTX4). One-way ANOVA with Tukey’s multiple comparison was used for quantification of mitophagy in cells and Two-way ANOVA with uncorrected Fisher’s LSD test for OXPHOS protein and mRNA expression quantification (except for isotope tracing which used Šídák’s multiple comparison test). Multiple comparison t-test was used for line graphs. Area over the curve (AOC) for IPITT and area under the curve (AUC) for IPGTT, RER and energy expenditure used unpaired two-tailed Student’s *t*-test. Graphs illustrate individual values, means and standard deviation (SD) unless otherwise indicated with statistical significance of p<0.05 of minimum n =3 animals or independent passages per group.

## 3 Results

### 3.1 Skeletal Muscle-Specific STX4 Ablation Inherently Induces Insulin Resistance

We generated skmSTX4-iKO mice using a skeletal muscle-specific gene knockout model as previously used [8,19] by crossing Dox-inducible *HSA-rtTA/TRE-Cre^+/-^* and *Stx4^fl/fl^* construct lines to excise the mouse *Stx4* gene. Compared to Dox-treated non-inducible single *Stx4^fl/fl^;Cre^-/-^* (CTRL) mice, we observed significant depletion in STX4 protein abundance in soleus (oxidative fiber rich), TA (glycolytic fiber rich) and GAS (mixed fibers) skeletal muscles (Figure 1a) with no significant change in non-skeletal muscle tissues (heart and liver) (Figure 1b). *Stx4* mRNA levels across these muscle groups were also decreased (Figure S1a-c). Significantly reduced gains in bodyweight (Figure 1c) and weights of large muscle groups such as quadriceps and GAS for the skmSTX4-iKO male mice were also detected (Table S3). Furthermore, IPITT (Figure 1d) and IPGTT (Figure S1d) revealed significantly decreased insulin sensitivity (Figure 1d) and impaired glucose tolerance (Figure S1d) in non-obese skmSTX4-iKO male mice compared to age-matched CTRL. Female skmSTX4-iKO mice showed similar changes (Figure S2, Table S4). These data indicate that inducible STX4 ablation selectively in skeletal muscle leads to the development of insulin resistance.

**Figure 1.**
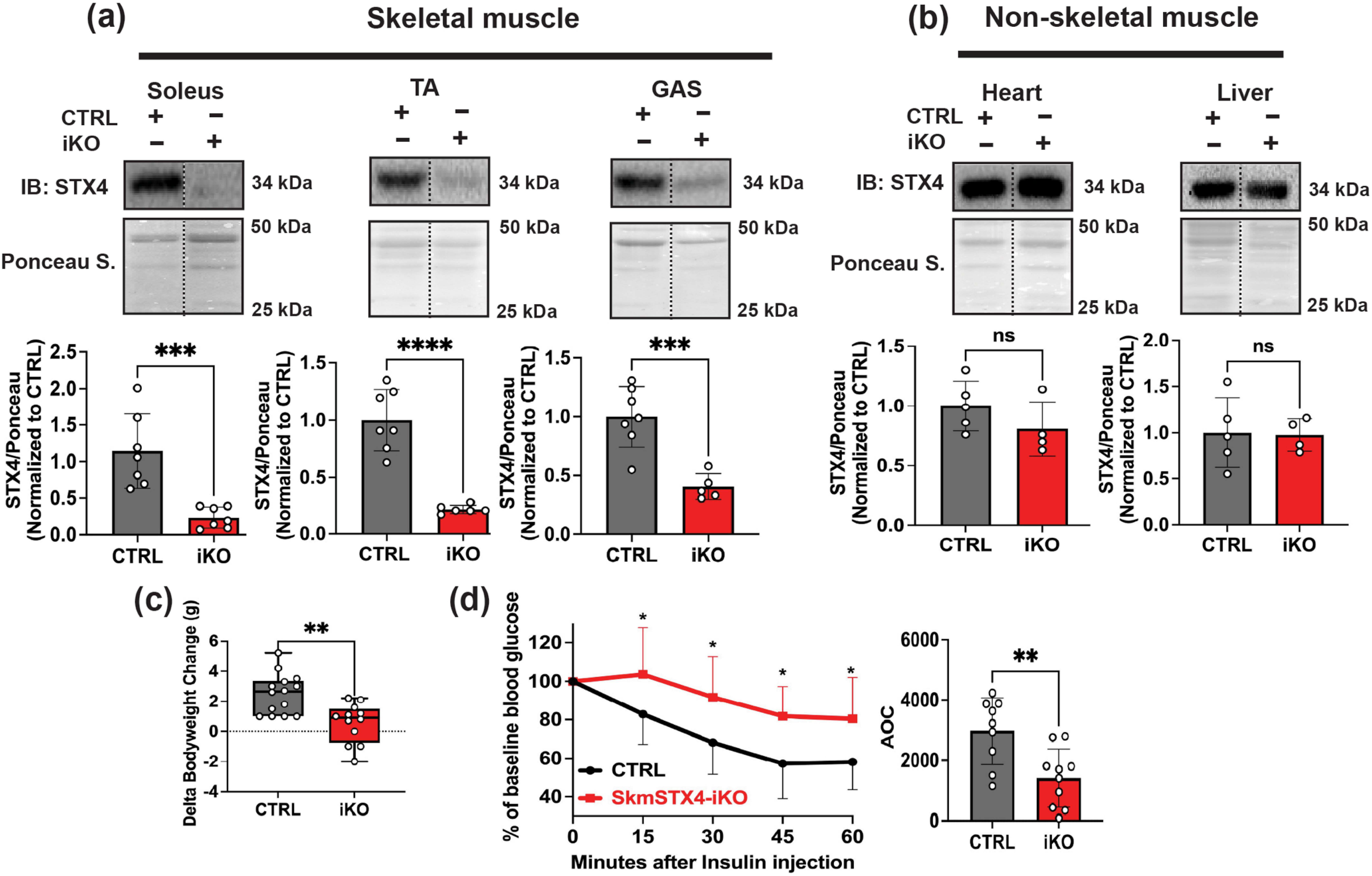
SkmSTX4-iKO male mice develop inherent insulin resistance. **(a)** Representative immunoblot (top) and quantification (bottom) of STX4 protein abundance across skeletal muscle depots including soleus, Tibialis anterior (TA) and gastrocnemius (GAS) alongside **(b)** non-skeletal muscle organs of heart and liver from control (CTRL, Grey) and skmSTX4-iKO (iKO, Red) 20-week-old male mice normalized against total protein lysate stained with ponceau (n=4-7 mice/group). **(c)** Delta bodyweight changes of initial weight to final weight of control (CTRL, Grey) and skmSTX4-iKO (iKO, Red) 20-week-old male mice (n=12-14 mice/group). **(d)** Intraperitoneal insulin tolerance test (IPITT) of control (CTRL, Black/Grey) and skmSTX4-iKO (iKO, Red) 20-week-old male mice with area over the curve (AOC) bar chart (n=9-10 mice/group). Data in bar and line graphs shown as mean ± SD. Statistical significance was determined by unpaired two tailed Student’s t test **(a, b, c** and **d** (AOC)) and multiple t-test model (**d**). **p<0.01, ***p<0.001, ****p<0.0001, ns = not significant. Vertical dashed lines indicate the splicing of lanes within the same gel exposure. IB: Immunoblot, Ponceau S.: Ponceau stain.

### 3.2 Skeletal Muscle-Specific STX4 Ablation Decreases Whole Body Energy Homeostasis and Muscle Performance via Impaired Mitochondrial Function

To assess the impact of skeletal muscle specific STX4 ablation on whole body metabolism in skmSTX4-iKO mice, we initially acclimatized CTRL and skmSTX4-iKO male mice in metabolic caging units overnight and then captured data every hour for up to 72 hours. The RER was significantly lower in skmSTX4-iKO male mice (Figure 2a), suggesting a shift towards fat as a predominant metabolic fuel [8]. Interestingly, we observed a significant decrease in energy expenditure in the skmSTX4-iKO male mice compared to their CTRL counterparts (Figure 2b). However, our metabolic cage measurements showed no significant differences in total food and water consumption (Figure S3). Based on previous studies in adipose tissue [S6] the decreased RER (Figure 2a) and energy expenditure (Figure 2b) could be a consequence of decreased mitochondrial activity. Thus, we next assessed mitochondrial function via OCR in isolated primary FDB myofibers from CTRL and skmSTX4-iKO mice [8]. Notably, we found a significant reduction of OCR in the skmSTX4-iKO myofibers that flatlines across the basal and maximal respiration OCR phases compared to the CTRL (Figure 2c). Next, we assessed whether the skeletal muscle mitochondrial dysfunction presented in skmSTX4-iKO male mice was associated with weakened physical performance i.e. muscle strength as observed in conditions like aging [S7] and diabetes [S8]. Thus, we measured forelimb grip strength and found that skmSTX4-iKO male mice presented ∼25% reduced grip strength as compared to their CTRL counterparts (Figure 2d). These findings suggest that STX4 ablation in skeletal muscle impedes whole body energy efficiency and physical performance through impaired mitochondrial function.

**Figure 2.**
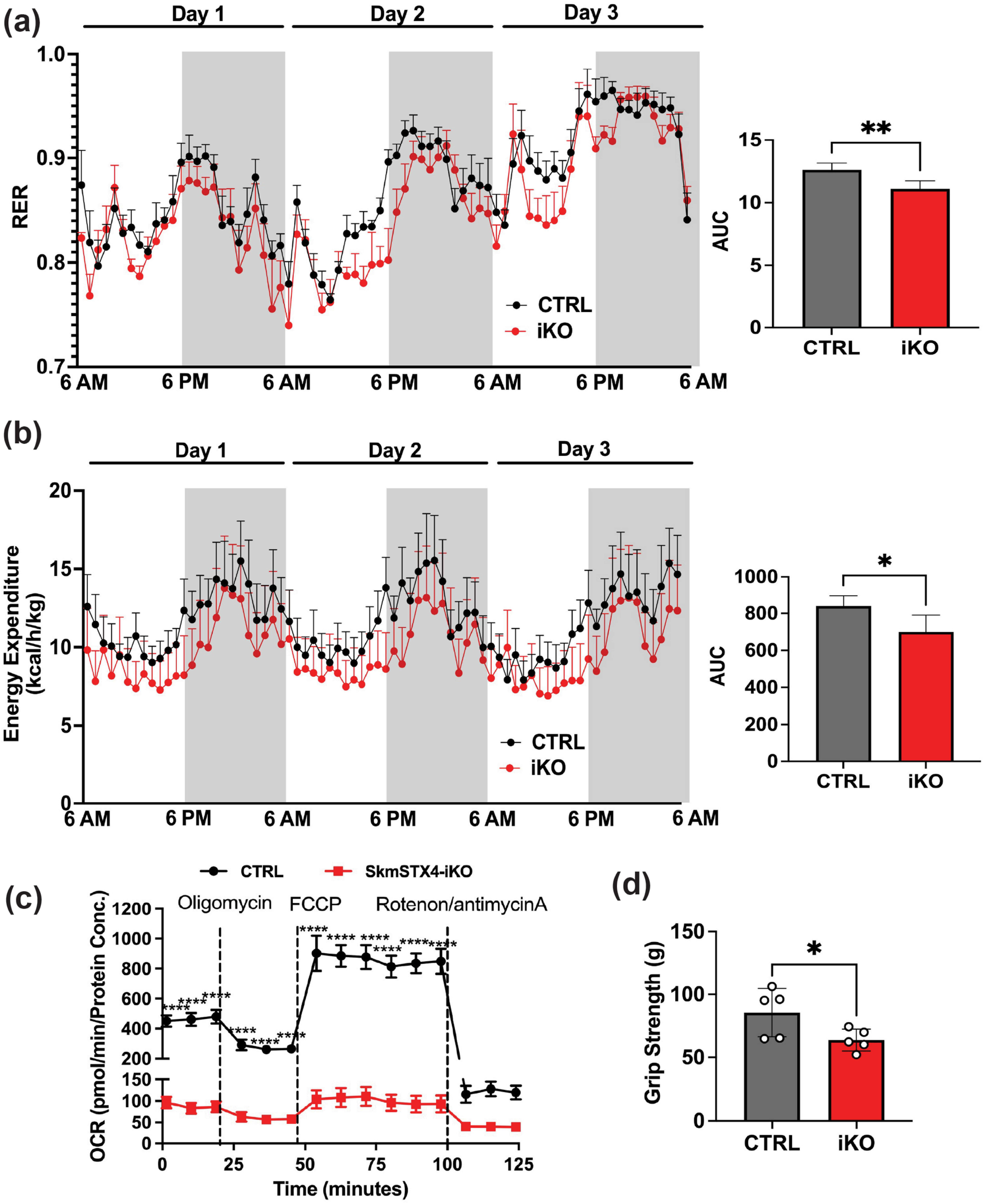
SkmSTX4-iKO male mice develop impaired whole-body metabolism and muscle function via impaired mitochondrial activity. **(a-b)** Metabolic caging analyses of the control (CTRL, Grey/Black) and skmSTX4-iKO (iKO, Red) 26-week-old male mice collected over a 72-hour period. Left: **(a)** Respiratory exchange ratio (RER) and **(b)** Energy expenditure. Data presented as line charts with white shading representing daytime (inactive) phases (6 am – 6 pm) and grey shading representing nighttime (active) phases (6 pm – 6 am). Right: Corresponding area under the curve (AUC) calculation (n=6-10 mice/group). **(c)** Extracellular flux analysis (Seahorse) to assess the mitochondrial oxygen consumption rate (OCR) of isolated Flexor digitorium brevis myofibers from control (CTRL, Black) and skmSTX4-iKO (iKO, Red) 20-week-old male mice (n=5-10 mice/group). **(d)** Forelimb grip strength between control (CTRL, Grey) and skmSTX4-iKO (iKO, Red) 20-week-old male mice (n=5 mice/group). Data in bar and line graphs shown as mean ± SD. Statistical significance was determined by unpaired two tailed Student’s t test **(a** (AUC)**, b** (AUC), **d**) and mixed effects analysis with Šídák’s multiple comparison test (**c**). *p<0.05, **p<0.01, ****p<0.0001.

### 3.3 STX4 Ablation Differentially Impacts Mitochondrial Structural Damage and ETC Abundances in Oxidative versus Glycolytic Rich Skeletal Muscles

As STX4 is located at both the sarcolemma and OMM in skeletal muscle [8] and its loss negatively impacts mitochondrial function, we next tested whether STX4 ablation impacts changes in skeletal muscle mitochondrial structure. To assess mitochondrial structure, we analyzed mitochondria in CTRL and skmSTX4-iKO mouse soleus (oxidative slow-twitch) and TA (glycolytic fast-twitch) by TEM, as previously described [8]. In soleus of skmSTX4-iKO male mice, smaller and more fragmented mitochondria were observed, with reduced mitochondrial area, as compared to CTRL (Figure 3a). In TA of skmSTX4-iKO male mice, a larger mitochondrial area was attributed to a bloated swollen damaged mitochondrial structure with abnormal cristae, as compared to CTRL (Figure 3b). Despite being non-obese, these skmSTX4-iKO mice presented mitochondrial damage reminiscent of those observed in HFD-exposed insulin resistant obese mice [8]. Given the differences observed in mitochondrial damage between the oxidative rich (soleus) (Figure 3a) and glycolytic rich (TA) (Figure 3b) skeletal muscle in skmSTX4-iKO male mice, we next evaluated changes in electron transport chain (ETC) complexes between these two different muscle types. We observed a significant decrease in ETC complexes I, II, and IV in oxidative fiber rich skmSTX4-iKO soleus muscle compared to CTRL mice (Figure 3c). In contrast, ETC complexes in the glycolytic fiber rich skmSTX4-iKO TA muscle were unchanged (Figure 3d), suggesting that STX4 ablation has a greater impact in ETC protein abundance in oxidative fiber types. We found similar mitochondrial phenotype and ETC heterogeneity between soleus and TA muscle in female skmSTX4-iKO mice (Figure S4). These data demonstrate that STX4 depletion impairs skeletal muscle mitochondrial structure, supporting that not only is STX4 necessary for the mitochondria in skeletal muscle, but also displays differential impacts on oxidative vs glycolytic muscles.

**Figure 3.**
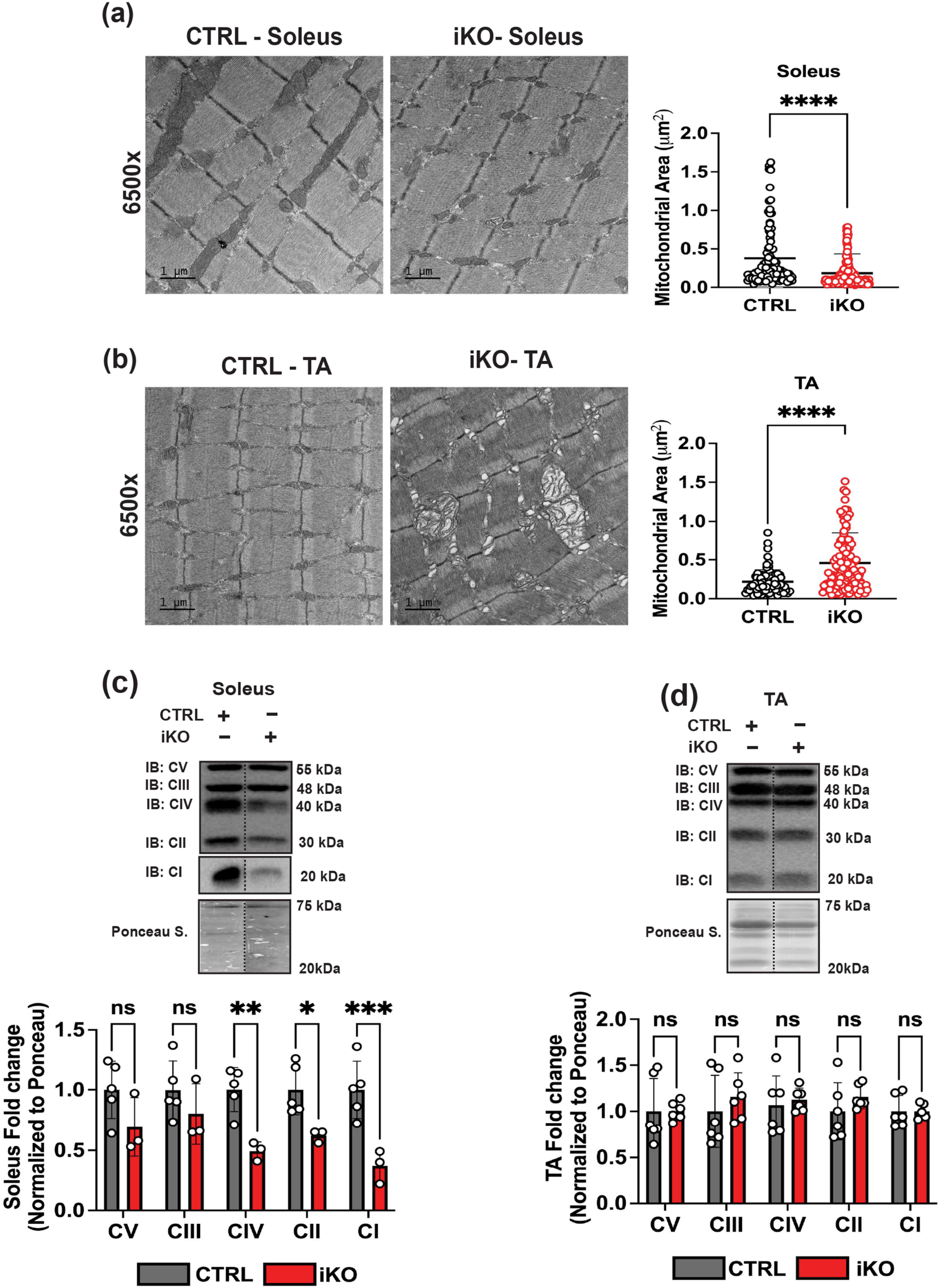
STX4 ablation has a heterogenous impact on mitochondrial structure and electron transport chain complex abundance between oxidative rich soleus and glycolytic rich TA muscle in male skmSTX4-iKO mice. Representative transmission electron microscopy images and mitochondrial area (μm^2^) quantification of mitochondrial from **(a)** soleus (n = 115-171 total mitochondria) and **(b)** TA muscle (n = 76-132 total mitochondria) from control (CTRL, Grey) and skmSTX4-iKO (iKO, Red) 20-week-old male mice. Black bar = 1 μm (6500X). Representative immunoblot (top) and quantification (bottom) of total electron transport chain (ETC) complexes V (CV), III (CIII), IV (CIV), II (CII) and I (CI) normalized against total protein lysate stained with ponceau in **(c)** soleus and **(d)** TA muscle from control (CTRL, Grey) and skmSTX4-iKO (iKO, Red) 20-week-old male mice (n=3-6 mice/group). Data in bar graphs shown as mean ± SD. Statistical significance was determined by unpaired two tailed Student’s *t* test (**a**, **b**) and two-way ANOVA with uncorrected Fisher’s LSD test (**c, d**). *p<0.05, **p<0.01, ***p<0.001, ****p<0.0001, ns = not significant. Vertical dashed lines indicate the splicing of lanes within the same gel exposure. IB: Immunoblot, Ponceau S.: Ponceau stain.

### 3.4 STX4 Ablation Does Not Impact Mitochondrial TCA Cycle for Glucose Metabolism

With the damaged mitochondrial phenotype in STX4 depleted muscle, we next assessed citrate synthase activity, a marker of the TCA cycle in glucose metabolism [8]. However, we observed no significant difference in the GAS muscle between CTRL and skmSTX4-iKO male mice (Figure 4a). With TCA cycle being a dynamic process, we next utilized a live culture system of the rat L6.GLUT4myc myotube cell line as successfully used in previous studies on mitochondrial activity [8,9]. Following a paradigm of 96 hours siRNA transfection, we found that siSTX4 L6.GLUT4myc myotubes showed significant decreases in ETC I, II and IV complexes compared to siCON (Figures 4b-c). With decreases in these ETC complexes, we next assessed the impact of STX4 ablation on the TCA cycle for glucose metabolism via incubation with [U-^13^C]glucose labelled media, as previously performed [8]. However, we observed no significant difference in the levels of glycolytic (Figure 4d) or TCA (Figure 4e) metabolites between siSTX4 and siCON L6.GLUT4myc myotubes, suggesting that STX4 ablation has no impact on the TCA cycle for mitochondrial glucose metabolism in skeletal muscle. Similarly, we found STX4 ablation does not alter markers of lactate conversion (glycolysis), fatty acid oxidation or canonical insulin signaling in skeletal muscle (Figure S5). Overall, our data suggests that the effect of STX4 ablation on inducing insulin resistance and mitochondrial dysfunction does not involve glycolytic and mitochondrial glucose metabolism in skeletal muscle.

**Figure 4.**
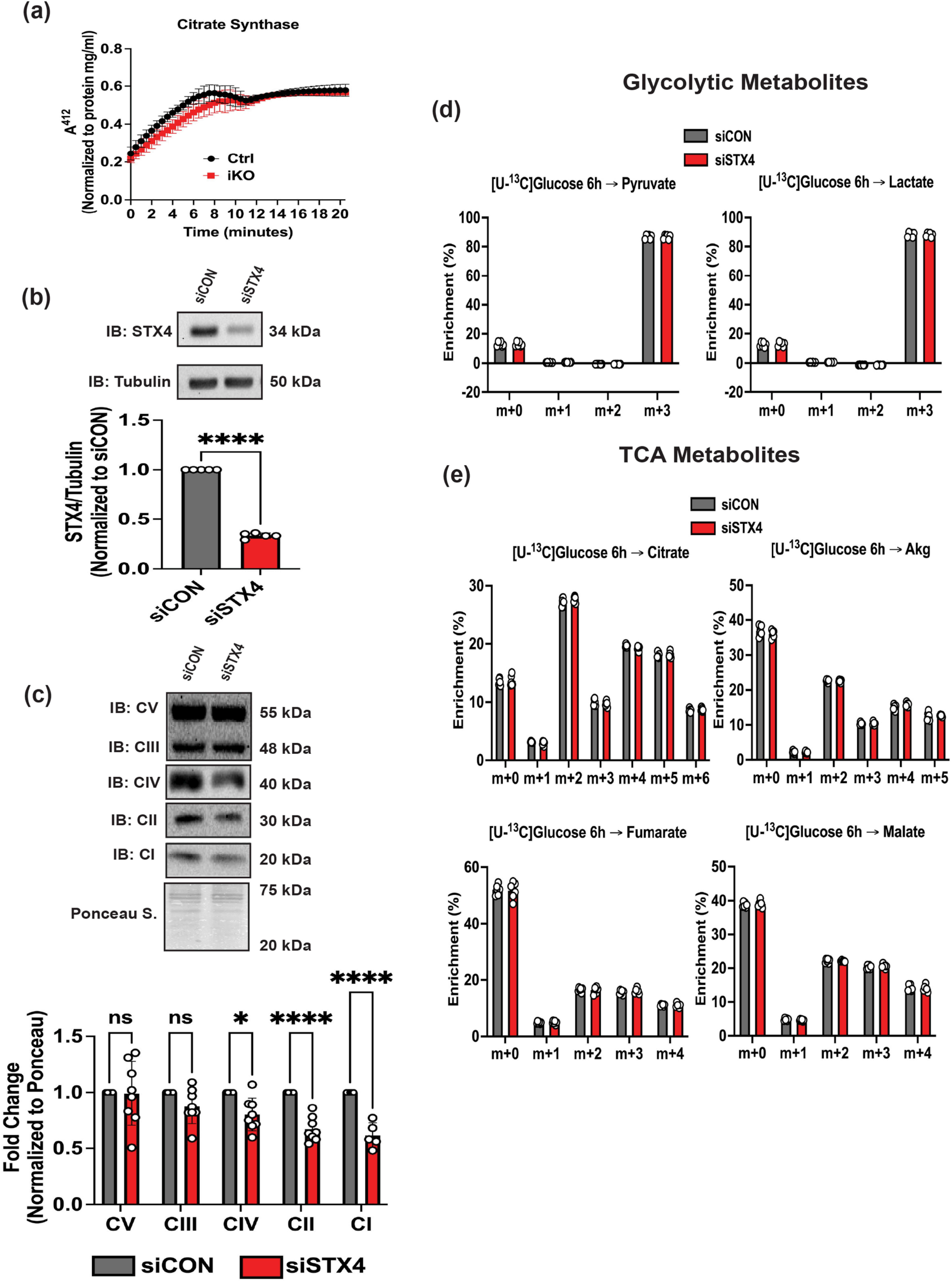
STX4 ablation in skeletal muscle does not impact mitochondrial glucose metabolism. (**a**) Citrate synthase activity in the gastrocnemius muscle from control (CTRL, Black) and skmSTX4-iKO (iKO, Red) 20-week-old male mice (n=3 mice/group). (**b**) Representative immunoblot (top) and quantification (bottom) of STX4 protein abundance normalized against Tubulin (n=5 independent cell passages). (**c)** Representative immunoblot (top) and quantification (bottom) of total electron transport chain (ETC) proteins complexes V (CV), III (CIII), IV (CIV), II (CII) and I (CI) normalized against total protein lysate stained with ponceau (n=5-8 independent cell passages). (**c**) Schematic and quantification of (**d**) glycolytic metabolites – Pyruvate and Lactate, and (**e**) tricarboxylic acid (TCA) metabolites – Citrate, Akg, Fumarate, and Malate post-[U-^13^C] glucose incubation for 6 hours (n=6 independent cell passages) with readings normalized to total protein levels in whole cell lysate. Data in bar and line graphs shown as mean ± SD. Statistical significance was determined using multiple unpaired t-tests (**a**), unpaired two tailed Student’s *t* test (**b**), two-way ANOVA with uncorrected Fisher’s LSD test (**c**) and Šídák’s multiple comparison test (**d-e**). *p<0.05, ***p<0.001, ****p<0.0001, ns = not significant. IB: Immunoblot, Ponceau S.: Ponceau stain.

### 3.5 STX4-Depletion Reduces Mitochondrial DNA Copy Number and Impacts Mitochondrial Biogenesis Regulators

We previously established that STX4 plays an important role in mitochondrial dynamics via its interaction with mitochondrial fission protein Drp1 [8]. Given the coordinated nature of MQC [5,6], STX4 may function in other mitochondrial processes, such as mitochondrial biogenesis. Thus, we assessed mtDNA copy number via qPCR of mtDNA levels and found significantly depleted mtDNA copy number in skmSTX4-iKO mouse muscle (Figure 5a). To determine whether this decrease in mtDNA copy number was attributed to impaired mitochondrial biogenesis, we next evaluated PGC1-α, a well-established master regulator of mitochondrial biogenesis and key mitochondrial biogenesis regulators NRF1 and TFAM [5–7]. Given the short half-life of PGC1-α [9] and NRF1 [S9-10] proteins, we assessed mitochondrial biogenesis genes mRNA levels and found a significant decrease of *Ppargc1a* (PGC1-α) and *Tfam* (Figures 5b) in oxidative soleus muscle from skmSTX4-iKO mice only, with no significant difference in *Nrf1* expression (Figure 5b). However, we found no significant difference of mitochondrial biogenesis genes in glycolytic TA muscle (Figure S6). Similarly, our siSTX4 L6.GLUT4myc myotubes presented significant decreases of mtDNA copy number (Figure 5c). Interestingly, the mRNA levels across all biogenesis genes (Figure 5d) and protein abundances for PGC1-α and NRF1 (Figures 5e-f) were reduced in siSTX4 L6.GLUT4myc myotubes, which could be attributed to the key role mitochondrial biogenesis plays in establishing the mitochondrial network in myotube differentiation [20,21]. Taken together, these findings suggest that STX4 ablation reduces mtDNA copy number and leads to mitochondrial biogenesis dysregulation.

**Figure 5.**
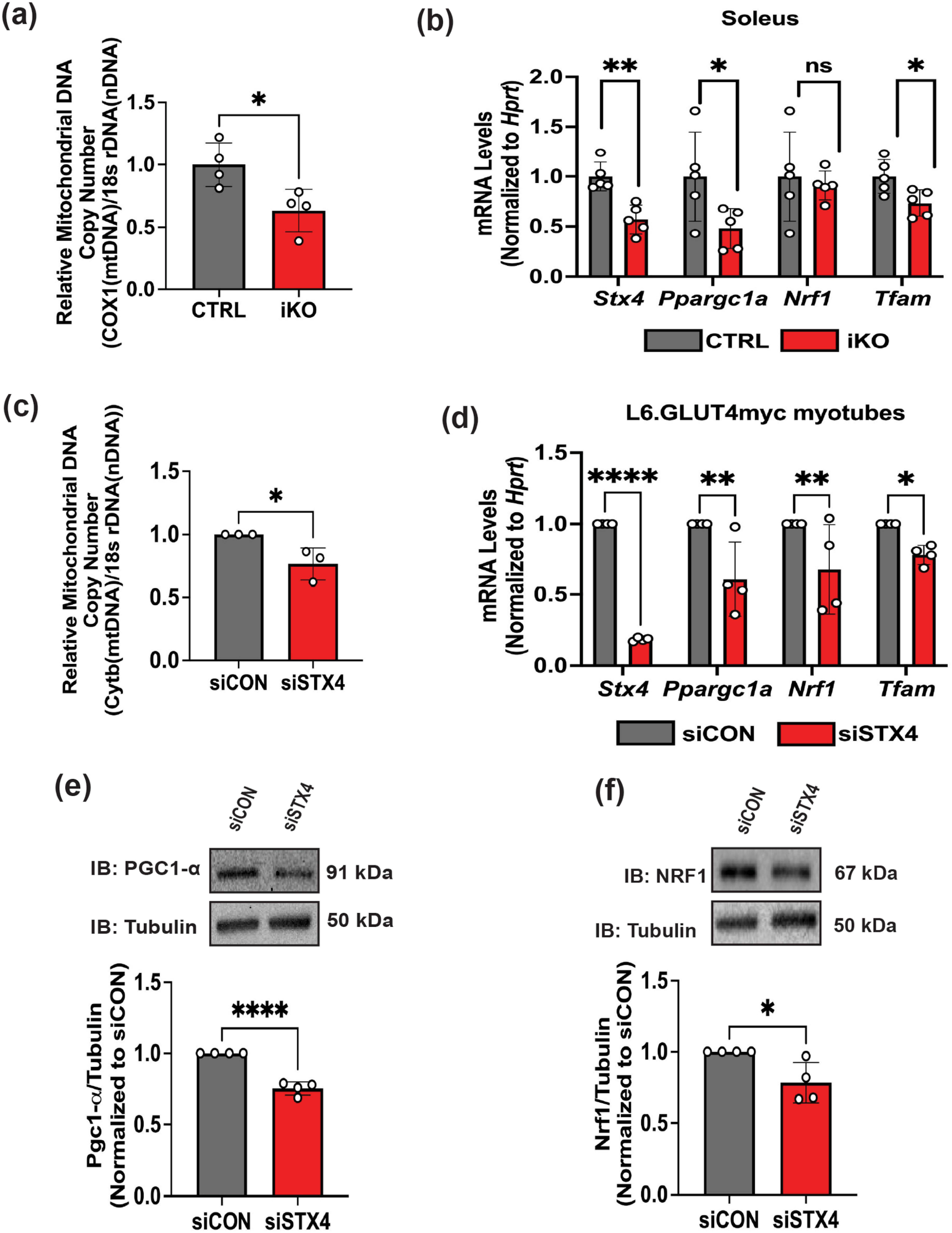
STX4 ablation in skeletal muscle impact mitochondrial DNA copy number and mitochondrial biogenesis. (**a**) Quantification of mitochondrial DNA (mtDNA) copy number between control (CTRL, Grey) and skmSTX4-iKO (iKO, Red) 20-week-old male mouse gastrocnemius muscle (n=4 mice/group), expressed as a ratio of *Cox1* gene (mtDNA) to *18s ribosomal DNA* (Nuclear DNA). (**b**) Quantification of mRNA expression for *Stx4*, *Ppargc1a* (PGC1-α)*, Nrf1* and *Tfam* in the soleus muscle between control (CTRL, Grey) and skmSTX4-iKO (iKO, Red) 20-weeks-old male mice normalized to *Hprt* housekeeping gene (n=5 mice/group). (**c**) Quantification of mtDNA copy number between siCON (Grey) and siSTX4 (Red) L6.GLUT4myc myotubes expressed as a ratio of *Cytochrome B* (*Cytb*) (mtDNA) to *18s ribosomal DNA* (nuclear DNA) (**d**) Quantification of mRNA expression for *Stx4, Ppargc1a* (PGC1-α)*,Nrf1* and *Tfam* between siCON (Grey) and siSTX4 (Red) L6.GLUT4myc myotubes, normalized to *Hprt* housekeeping gene (n=5 independent cell passages). Representative immunoblot (top) and quantification (bottom) of (**e**) PGC1-α and (**f**) NRF1 protein abundance normalized to Tubulin between siCON (Grey) and siSTX4 (Red) L6.GLUT4myc myotubes (n=4 independent cell passages). Data in bar graphs shown as mean ± SD. Statistical significance was determined using unpaired two tailed Student’s *t* test (**a**, **c**, **e** and **f**) and two-way ANOVA with uncorrected Fisher’s LSD test (**b**, **d**). *p<0.05, **p<0.01, ****p<0.0001, ns = not significant. IB: Immunoblot.

### 3.6 STX4 ablation impairs mitochondrial turnover via canonical mitophagy

Mitochondrial biogenesis is interlinked with mitochondrial turnover by mitophagy [5–7]. Thus, we next evaluated the requirement for STX4 in mitophagy. Being a dynamic process, we initially assessed mitophagy using live cell imaging by dual transfection of siCON or siSTX4 L6.GLUT4myc myoblasts with mt-Keima plasmid, a mitochondrial targeting dual excitation protein to distinguish non-lysosomal-associated mitochondria from lysosomal-associated mitochondria undergoing mitophagy (Figure 6a) [S5]. Although we observed no significant difference in mt-Keima red signal under basal conditions, STX4 knockdown significantly reduced mt-Keima red signal under CCCP exposure (a mitophagy inducer) [S4-5], consistent with a tempered mitophagic response (Figures 6b-c). To confirm that the reduced mt-Keima red was due to impaired mitochondria-lysosomal co-localization, we used MitoTracker (red) and Lysotracker (green) reagents to mark both organelles and evaluated their colocalization (yellow) (Figure 6d). We observed reduced co-localization in siSTX4 10 μM CCCP-exposed L6.GLUT4myc myotubes (Figures 6e-f) versus control, suggesting that STX4 ablation in skeletal muscle impairs mitochondrial turnover by mitophagy. With one recent report suggesting a predominance of canonical PINK1-PARKIN mitophagy in oxidative-rich skeletal muscle [11], we assessed these proteins in both the soleus and TA muscle from skmSTX4-iKO male mice. Interestingly, we showed reduced PARKIN (Figure 7a) and PINK1 (Figure 7b) protein abundance in both skmSTX4-iKO soleus and TA muscle, suggesting that canonical mitophagy is impacted by STX4 ablation in both oxidative and glycolytic rich muscle. Similarly, we observed reduced PARKIN (Figure 7c) and PINK1 (Figure 7d) protein abundance in siSTX4 L6.GLUT4myc myotubes. Altogether, our data suggests that STX4 could be required for mitochondrial turnover by mitophagy in skeletal muscle and that this is associated with the canonical PINK1-PARKIN pathway.

**Figure 6.**
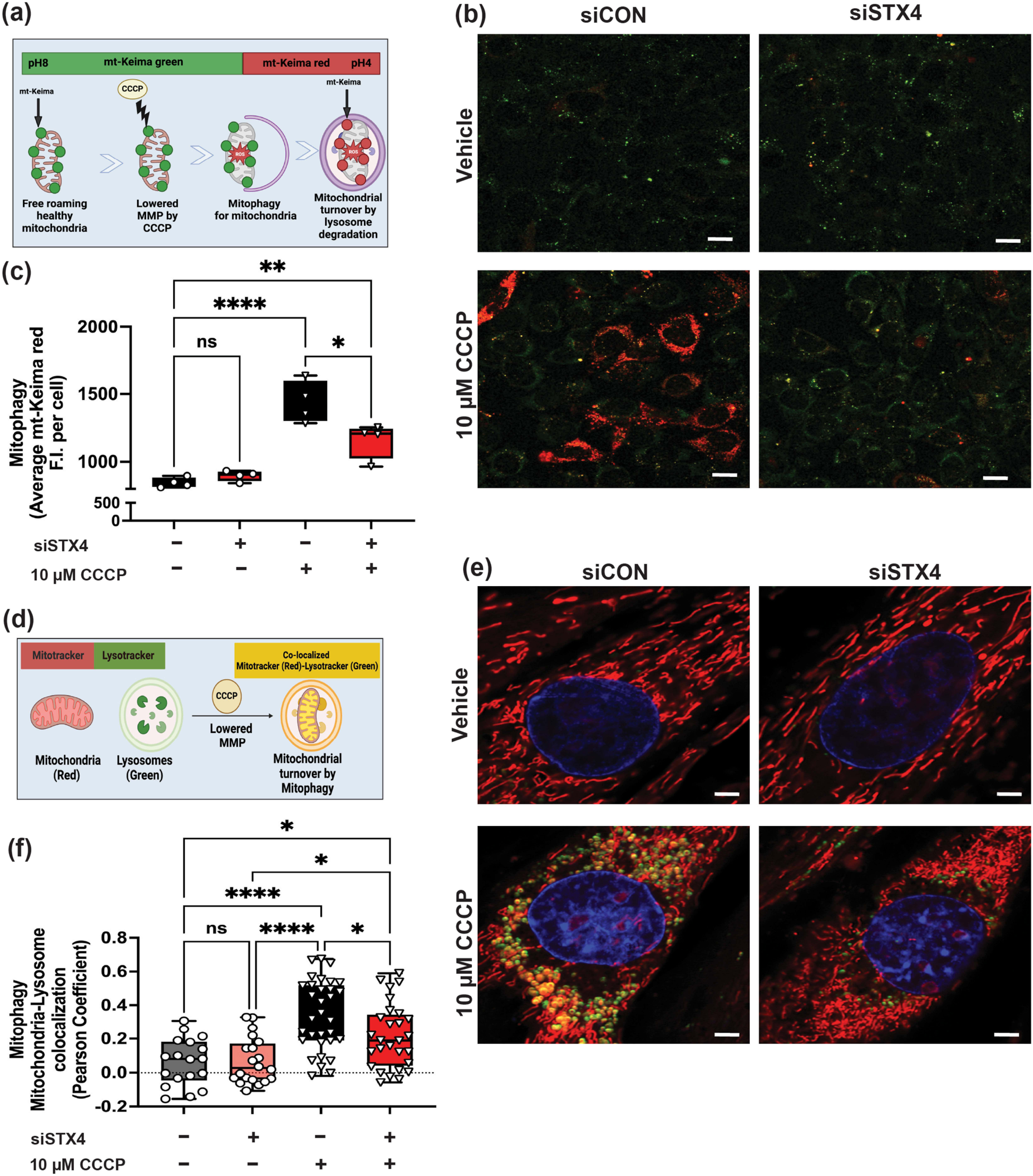
STX4 ablation in L6.GLUT4myc myoblasts and myotubes impairs mitochondrial turnover by mitophagy. (**a**) Schematic showing mt-Keima detection representative of active mitophagy. (**b-c**) Representative confocal fluorescence microscope image (**b**) and quantification by mean fluorescent intensity (F.I) (**c**) of active mitophagy (Red) across vehicle or 10 μM CCCP-treated siCON (Black/Grey) and siSTX4 (Red) L6.GLUT4myc myoblasts (n=3-4 independent cell passages). Scale bar represents 20 μm. (**d**) Schematic showing detection of mitochondria and lysosome co-localization representing active mitophagy. (**e-f**) Representative confocal fluorescence microscope image (**e**) and quantification of active mitochondria-lysosome colocalization (**f**) across vehicle or 10 μM CCCP-treated siCON (Black/Grey) and siSTX4 (Red) L6.GLUT4myc myotubes (n=3 independent cell passages). Scale bar represents: 10 μm. Box plot represents the mean ± SD (**c, f**). Statistical significance was determined using one-way ANOVA with Tukey’s multiple comparison test (**c, f**). *p<0.05, **p<0.01, ****p<0.0001, ns = not significant. Images created using BioRender.com (**a, d**).

**Figure 7.**
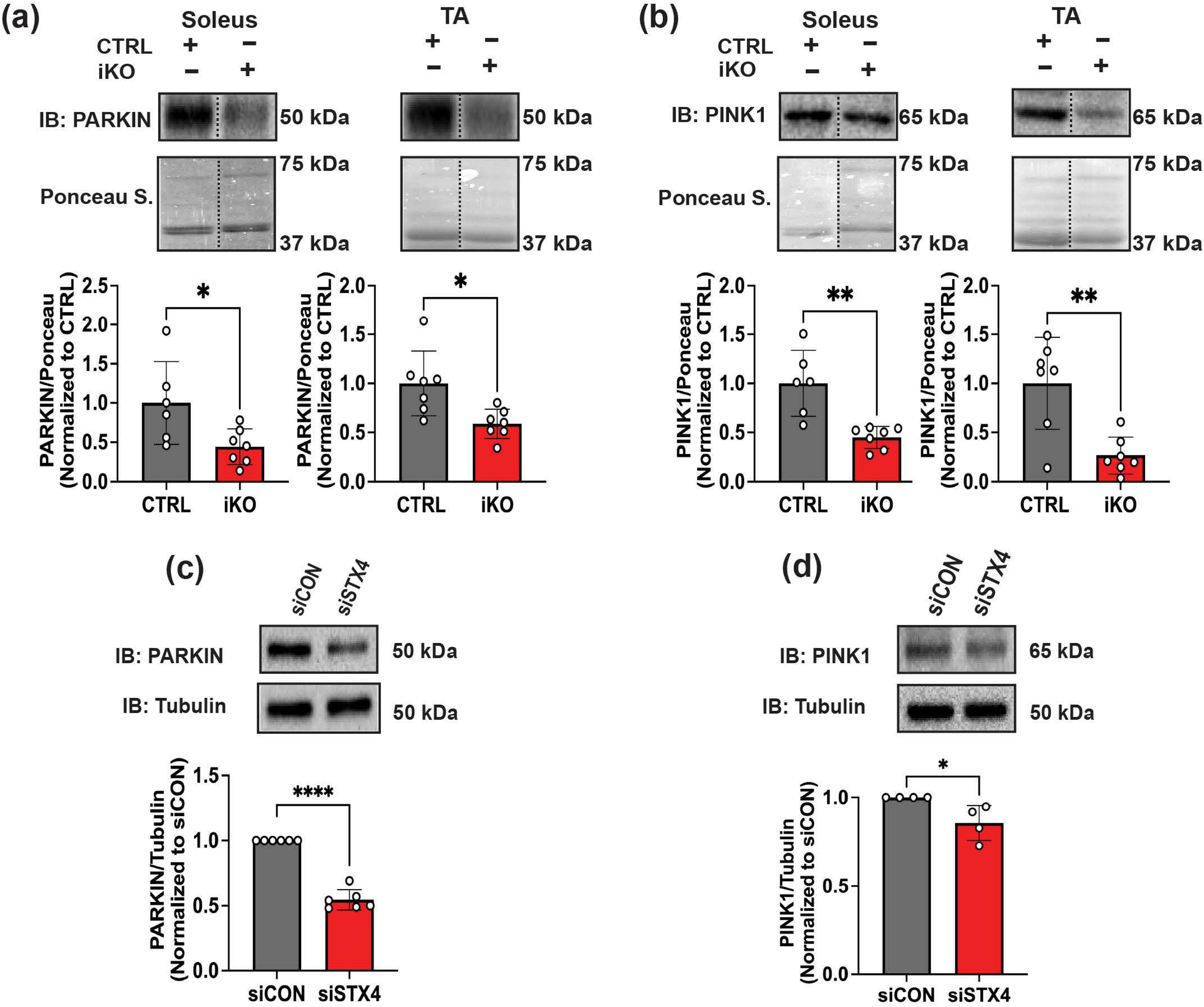
STX4 ablation impairs PINK1-PARKIN canonical mitophagy in skeletal muscle. Representative immunoblot (top) and quantification (bottom) of (**a**) PARKIN and (**b**) PINK1 protein abundance in control (CTRL, Grey) and skmSTX4-iKO (iKO, Red) 20-week-old male mice soleus and TA muscle normalized to total protein lysate by ponceau staining (n=6-7 mice/group). Representative immunoblot (top) and quantification (bottom) of (**c**) Parkin and (**d**) PINK1 protein abundance normalized to Tubulin in siCON (Grey) and siSTX4 (Red) L6.GLUT4myc myotubes (n=3-6 independent cell passages). Data in bar graphs shown as mean ± SD. Statistical significance was determined by unpaired two tailed Student’s *t* test (**a-d**). *p<0.05, **p<0.01, ****p<0.0001. Vertical dashed lines indicate the splicing of lanes within the same gel exposure. IB: Immunoblot, Ponceau S.: Ponceau stain.

## 4 Discussion

This study revealed that STX4 ablation in non-obese skeletal muscle induced insulin resistance and a mitochondrial phenotype reminiscent of skeletal muscle in diabetogenic obese C57BL6/J male mice [8], as demonstrated by damaged mitochondrial structure and impaired OCR contributing to dysregulated whole body metabolism. Furthermore, we explored the role of STX4 in MQC beyond mitochondrial dynamics [8] and found that STX4 ablation impaired both mitochondrial biogenesis and mitochondrial turnover by mitophagy. Taken together, these findings suggest that STX4 is indispensable for mitochondrial homeostasis and MQC in skeletal muscle.

Our previous investigations revealed no changes in ETC abundance within STX4 enriched whole hindlimb muscle [8]. However, due to the larger proportion of glycolytic rich muscle mass (e.g. GAS and TA) compared to oxidative (e.g. soleus) in whole hindlimb, our skmSTX4-iKO mice revealed fiber specific divergences in mitochondrial structure/area, damage and ETC abundance. In particular, oxidative soleus presented a fragmented mitochondrial phenotype with reduced abundances of ETC I, II, and IV, whilst glycolytic TA presented a swollen bloated mitochondrial phenotype with no change in ETC abundance in skmSTX4-iKO mice. Similarly, L6.GLUT4myc myotubes, a fast-twitch origin cell line with reported greater oxidative capacity, presented reductions of ETC complexes I, II and IV upon STX4 ablation [22,23]. Indeed, STX4 ablation in brown adipose tissue (BAT), which possesses high mitochondrial density and oxidative capacity, also resulted in mitochondrial dysfunction, smaller mitochondrial size and reduced abundances across all ETC complexes [24]. On the other hand, knockout of the *Survival of motor neuron 1* (*SMN1*) gene in TA muscle from C57BL6 mice presented mitochondrial phenotype reminiscent to those observed in our skmSTX4-iKO mice [25]. Our observed differences of STX4 ablation on mitochondrial damage between oxidative vs glycolytic fibers could be attributed to the innate mitochondrial homeostasis differences between these fibers [26]. For example, in the BAT study, STX4 in mitochondrial homeostasis was attributed to its interactive role in UCP1 protein stability [24]. Although UCP1 is predominantly expressed in BAT [27], a known homolog expressed in striated muscle is UCP3, which shares 59% homology with UCP1 and is involved in respiratory regulation, is present in higher abundance in glycolytic muscle than oxidative [27,28]. Although UCP3 is one of many potential unknown candidates for STX4 in skeletal muscle mitochondria, the heterogeneity of metabolically distinct multinucleated fiber types in skmSTX4-iKO mice compared to the BAT-STX4-iKO mice [2,8,24] requires future in-depth investigation on the single fiber level (oxidative vs glycolytic). In particular, technologies such as single fiber and spatial proteomics [29,30] in skmSTX4-iKO muscle would be useful in discerning STX4’s fiber specific role(s) in mitochondrial regulation.

Previously, we determined that the STX4 enrichment improved mitochondrial function in HFD-exposed male mice, leading to efficiency at expending energy with increased activity coupled with lower energy expenditure [8]. On the other hand, the skeletal muscle mitochondrial dysfunction in skmSTX4-iKO male mice was associated with reduced energy expenditure albeit with no significant difference in food and water consumption compared to CTRL counterparts. Previous reports indicate that mitochondrial dysfunction could be a contributing factor for reduced energy expenditure [S6] [31]. In particular, reduced energy expenditure and muscle mitochondrial dysfunction are associated with exercise intolerance [31], which in T2D [32] diminishes the ameliorative effects of exercise to improve insulin sensitivity. Although it remains to be determined whether STX4 deficiency is a factor in exercise intolerance, future studies could investigate the requirements and/or benefits of skeletal muscle STX4 in exercise.

To the best of our knowledge, this is the first study that has associated the ablation of a Syntaxin protein with the downregulation of mitochondrial biogenesis regulators. We reported previously that skeletal muscle specific STX4 enrichment boosted mtDNA copy number, postulating that this effect could be attributable to enlargement of mitochondrial size via Drp1 pS637 inactivation [8]. However, studies into Drp1 on mtDNA copy number and mitochondrial biogenesis in skeletal muscle have been unclear. Indeed, homozygous [33] but not heterozygous skeletal muscle-specific Drp1 knockout mice [34] displayed increase in mtDNA copy number with upregulation of PGC1-α, whilst Drp1 enrichment does not affect mitochondrial biogenesis or DNA copy number [35]. Similarly, STX17, a Syntaxin protein that like STX4 is linked with mitochondrial dynamics and mitophagy reported little impact on mtDNA copy number [36], suggesting that role in modulating mitochondrial biogenesis is specific to STX4. Thus, it remains to be determined whether the association of STX4 ablation with mitochondrial biogenesis was causal or consequential, and future studies into novel STX4 mitochondrial interacting proteins could bridge this gap.

Although mitochondrial biogenesis and mitophagy have counter opposing activities in determining the abundance of mitochondria [5–7], our paradoxical results revealed that STX4 ablation negatively affects both processes. Interestingly, the paradoxical pattern of coordination between mitochondrial biogenesis and mitophagy simultaneously was also found in neuronal [37] and adipose [38] tissue studies for PGC1-α and PARKIN, proteins both significantly reduced in siSTX4 L6.GLUT4myc myotubes. Prior studies showed that PGC1-α and PARKIN can synergistically interact to enable protein stability between each other. Mitochondrial studies in BAT have shown that STX4 stabilizes mitochondrial proteins such as UCP-1 [24]. Thus, future studies may focus on whether STX4 stabilizes the regulatory proteins involved in mitochondrial biogenesis and mitophagy.

Although the prior consensus regarding mitophagy in diabetes was that elevations in mitophagy markers like PINK1-PARKIN in skeletal muscle were associated with the onset of T2D [39], recent studies suggests increased mitophagy actually serves as a protective response for the onset of insulin resistance [10,39]. However, as insulin resistance progresses from prediabetes to T2D, muscle biopsies have shown downregulation of PINK1-PARKIN alongside mitochondrial dysfunction [10,16,17]. Likewise, aging studies also demonstrated mitochondrial and MQC dysfunction in muscle [S7]. With skmSTX4-iKO mice presenting similar mitochondrial and MQC dysfunction patterns in skeletal muscle, it remains to be determined whether muscle-specific STX4 enrichment could evoke restorative effects and enhance canonical mitophagy to efficiently remove damaged mitochondria in diabetogenic and aging muscle. Evidence supporting this notion can be found in our global STX4 enriched mice, which presented the highest life expectancy recorded for an aging transgenic mouse model with maintained insulin sensitivity [40].

In summary, our study revealed that STX4 plays a critical role in the regulation of mitochondrial homeostasis in skeletal muscle, by modulating mitochondrial biogenesis and mitochondrial turnover by mitophagy. Thus, our work shows that STX4 is a critical component in skeletal muscle mitochondrial homeostasis providing support for therapeutic targeting of STX4 to improve mitochondrial health in metabolic disorders such as prediabetes and T2D. However, the precise mechanisms linking STX4 with regulation of mitochondria and MQC in skeletal muscle remains to be determined.

## Supporting information

Supplementary References

Raw Blots

## Acknowledgements

This study was supported by grants from the National Institutes of Health (DK067912, DK1129712 and DK102233 to D.C.T; R01DK137515 to K.M) and American Heart Association (25CDA1454374 to R.B). Research reported in this publication included work performed in the City of Hope Light Microscopy, Pathology, Comprehensive Metabolic Phenotyping, Integrated Mass Spectrometry Shared Resources and Animal Care Program Auxiliary Services supported by the National Cancer Institute of the National Institutes of Health (P30CA033572). We thank Dr. Zhuo Li, and Ricardo Zerda at City of Hope Electron Microscopy Core Facility for the help of electron microscopy. The authors thanks Dr. Chathurani S. Jayasena (City of Hope) for reviewing the manuscript and providing helpful feedback.

## Ethics Standards

The authors certify that the manuscript complies with the ethical guidelines for authorship and publishing in the Journal of Cachexia, Sarcopenia and Muscle. All animal studies have been approved by the appropriate ethics committee. The manuscript does not contain clinical studies or patient data.

