## Supplemental Tables and Figures for "STX4 is indispensable for mitochondrial homeostasis in skeletal muscle"

**Table of Contents:**

**Supplemental Table 1 ……………………………………………………………………………………......S2**
**Supplemental Table 2 ……………………………………………………………………………………..S3**
**Supplemental Table 3 ……………………………………………………………………………….. . S4**
**Supplemental Table 4 ……………………………………………………………………………………..S5**
**Supplemental Figure 1 ……………………………………………………………………………………S6**
**Supplemental Figure 2 …………………………………………………………………………………....S7**
**Supplemental Figure 3 ……………………………………………………………………………………S8**
**Supplemental Figure 4 ……………………………………………………………………………..…..S9**
**Supplemental Figure 5 …………………………………………………………………………………..S11**
**Supplemental Figure 6 …………………………………………………………………………………..S13**

### Supplemental Table 1. Antibody List

| Antibody | Vendor | Cat# | Size | Source | Dilution | RRID |
|---|---|---|---|---|---|---|
| STX4 | In-house generated | N/A | 34 kDa | Rabbit | 1:5000 | N/A |
| Tubulin | Sigma | T5168 | 50 kDa | Mouse | 1:10,000 | 086M4773V |
| PGC1-α | Novus Biologics | NBP3-08971 | 91 kDa | Rabbit | 1:1000 | 5500005011 |
| NRF1 | Abcam | Ab175932 | 68 kDa | Rabbit | 1:1000 | 1008037-22 |
| PINK1 | Proteintech | 23274-1-AP | 65 kDa | Rabbit | 1:1000 | 00146611 |
| PARKIN | Biolegend | SIG-39530 | 50 kDa | Mouse | 1:500 | B322798 |
| LDHB | Proteintech | 14824-1-AP | 35 kDa | Rabbit | 1:1000 | |
| CPT1B | Proteintech | 22170-1-AP | 75 kDa | Rabbit | 1:1000 | |
| AKT | Cell Signaling | 2920S | 68 kDa | Mouse | 1:2000 | |
| pS473 AKT | Cell Signaling | 4060S | 68 kDa | Rabbit | 1:2000 | |
| OXPHOS Cocktail (Rodent) | Abcam | Ab110413 | CV – 55 kDa<br>CIII – 48 kDa<br>CIV – 40 kDa<br>CII – 30 kDa<br>CI – 20 kDa | Mouse | 1:2000 | 2101039661 |

**Supplemental Table 2. Primer List**

| Species | Gene | Forward Primer 5’-3’ | Reverse Primer 5’-3’ |
|---|---|---|---|
| Mouse | *Stx4* | CAG TAT CCG AGA GCT CCA TG | CCG AGC TCA GGA TGT TCT TCT C |
| | *Ppargc1a* | GAA TCA AGC CAC TAC AGA CAC CG | CAT CCC TCT TGA GCC TTT CGT G |
| | *Nrf1* | GGC AAC AGT AGC CAC ATT GGC T | GTC TGG ATG GTC ATT TCA TCA CCG C |
| | *Tfam* | | |
| | *18S rDNA (Nuclear Gene)* | TGG CTC ATT AAA TCA GTT ATG GT | GTC GGC ATG TAT TAG CTC TAG |
| | *Cox1 (Mitochondrial Gene)* | ACC ATC ATT TCT CCT TCT CCT A | TAG ATT TCC GGC TAG AGG TG |
| | *Hprt* | AAG CCT AAG ATG AGC GCA AG | TTA CTA GGC AGA TGG CCA CA |

| | | | |
|---|---|---|---|
| Rat | *Stx4* | CAC GAT CCG TGA ACT CCA TG | CTG CTG AGC TCA GAA TGT TCT TTT C |
| | *Ppargc1a* | TGA ACT ACG GGA TGG CAA C | AAG AGC AAG AAG GCG ACA C |
| | *Nrf1* | GGC GCA GCA CCT TTG GAG AAT GTG | CAT CGA TGG TGA GAG GGG GCA GTT C |
| | *Tfam* | | |
| | *18S rDNA (Nuclear Gene)* | TAG AGG GAC AAG TGG CGT TC | CGC TGA GCC AGT CAG TGT |
| | *Mt-cyb (Mitochondrial Gene)* | TCC ACT TCA TCC TCC CAT TC | CTG CGT CGG AGT TTA ATC CT |
| | *Hprt* | GGT CCA TTC CTA TGA CTG TAG ATT TT | CAA TCA AGA CGT TCT TTC CAG TT |

**Supplemental Table 3. Tissue and body weights of chow-fed CTRL and skmSTX4-iKO (Dox-induced) male mice.**

| | CTRL (n = 13) | skmSTX4-iKO (n = 11) | p-value (two-tailed t-test) |
|---|---|---|---|
| Bodyweight (g) | 27.68 ± 1.21 | 24.05 ± 1.20 | <0.0001 (****) |
| Tissue (% Body weight) | | | |
| Kidney | 1.29 ± 0.03 | 1.54 ± 0.07 | 0.0057 (**) |
| Heart | 0.71 ± 0.03 | 0.82 ± 0.07 | 0.1406 |
| Spleen | 0.35 ± 0.03 | 0.43 ± 0.05 | 0.2066 |
| Pancreas | 0.80 ± 0.03 | 0.91 ± 0.04 | 0.056 |
| Fat | 1.93 ± 0.17 | 1.94 ± 0.18 | 0.5067 |
| Liver | 5.19 ± 0.70 | 5.98 ± 0.12 | <0.0001 (****) |
| Lungs | 0.87 ± 0.10 | 0.88 ± 0.05 | 0.2486 |
| Brain | 1.51 ± 0.05 | 1.75 ± 0.04 | 0.0032 (**) |
| Quadriceps Muscle | 1.43 ± 0.13 | 1.00 ± 0.80 | <0.0001 (****) |
| Gastrocnemius Muscle | 1.14 ± 0.15 | 0.76 ± 0.15 | <0.0001 (****) |
| Tibialis Muscle | 0.72 ± 0.16 | 0.75 ± 0.20 | 0.1858 |
| Soleus Muscle | 0.26 ± 0.10 | 0.35 ± 0.19 | 0.3960 |

Data represents the average ± SD of 20-week-old male mice. Significant differences detected using unpaired two-tailed t-test.

**Supplemental Table 4. Tissue and body weights of chow-fed CTRL and skmSTX4-iKO (Dox-induced) female mice.**

| | CTRL (n = 8) | skmSTX4-iKO (n = 8) | p-value (two-tailed t-test) |
|---|---|---|---|
| Bodyweight (g) | 21.75 ± 2.55 | 19.62 ± 1.85 | 0.0384 (*) |
| Tissue (% Body weight) | | | |
| Kidney | 1.14 ± 0.10 | 1.21 ± 0.14 | 0.3905 |
| Heart | 0.65 ± 0.09 | 0.62 ± 0.18 | 0.7278 |
| Spleen | 0.26 ± 0.07 | 0.38 ± 0.10 | 0.0549 |
| Pancreas | 0.90 ± 0.21 | 0.99 ± 0.18 | 0.4569 |
| Fat | 1.85 ± 1.21 | 0.99 ± 0.36 | 0.0995 |
| Liver | 5.07 ± 0.62 | 5.71 ± 0.70 | 0.1396 |
| Lungs | 0.71 ± 0.10 | 0.76 ± 0.05 | 0.4403 |
| Brain | 1.79 ± 0.21 | 2.10 ± 0.12 | 0.0153 (*) |
| Quadriceps Muscle | 1.21 ± 0.17 | 0.73 ± 0.14 | <0.0001 (****) |
| Gastrocnemius Muscle | 0.96 ± 0.15 | 0.59 ± 0.13 | <0.0001 (****) |
| Tibialis Muscle | 0.45 ± 0.05 | 0.39 ± 0.09 | 0.0998 |
| Soleus Muscle | 0.05 ± 0.03 | 0.06 ± 0.05 | 0.6763 |

Data represents the average ± SD of 20-week-old female mice. Significant differences detected using unpaired two-tailed t-test.

Supplemental Figure 1.

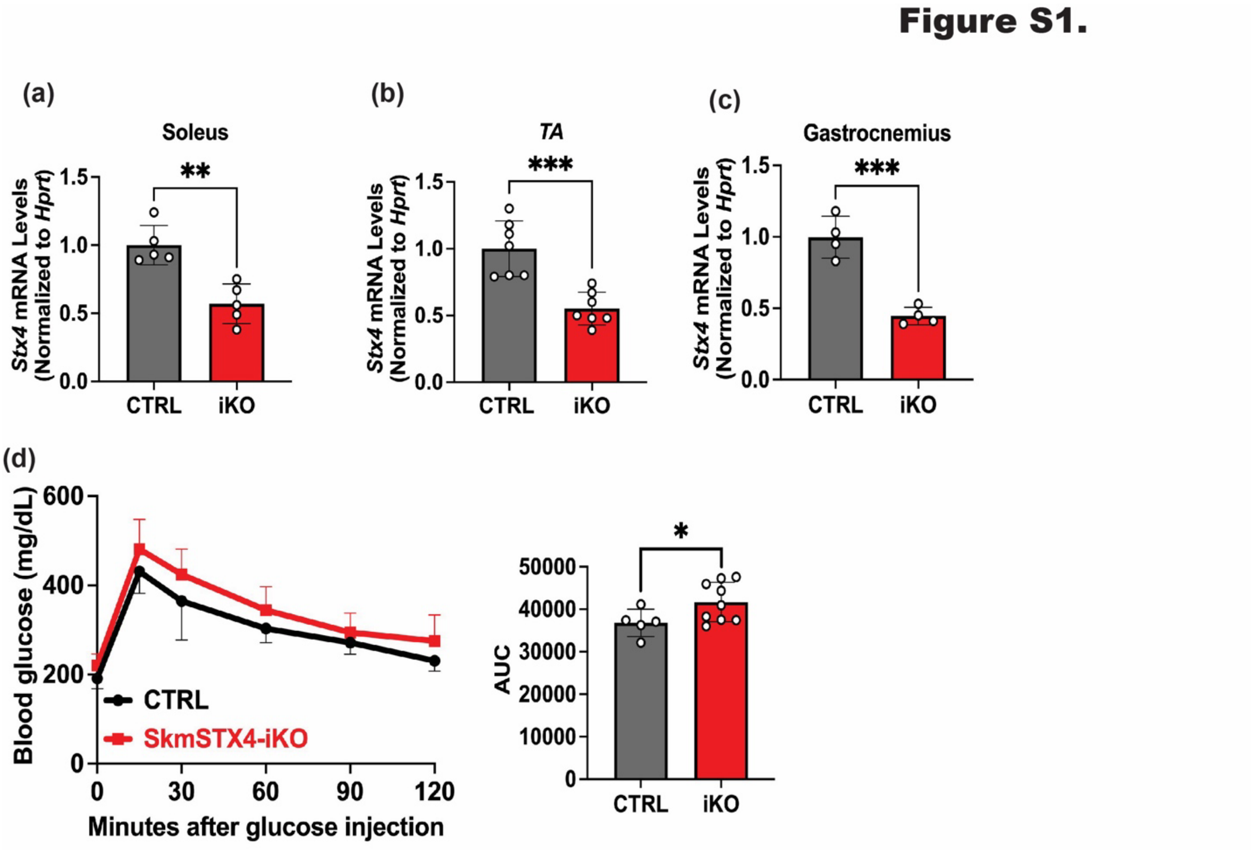


**Figure S1. Male skmSTX4-iKO mice present significantly reduced *Stx4* mRNA levels and impaired glucose tolerance.**

Quantification of mRNA levels for *Stx4* normalized against *Hprt* reference gene between control (CTRL, Grey) and skmSTX4-iKO (iKO, Red) 20-week-old male mice across (**a**) soleus, (**b**) Tibialis anterior (TA), and (**c**) gastrocnemius (GAS) muscle (n=4-6 mice/group). (**d**) Intraperitoneal glucose tolerance test (IPGTT) between control (CTRL, Black/Grey) and skmSTX4-iKO (iKO, Red) 20-week-old male mice with area under the curve (AUC) bar chart (n=5-9 mice/group). Data in bar and line graphs shown as mean ± SD. Statistical significance was determined by unpaired two tailed Student's t test (**a, b, c** and **d** (AOC)) and multiple t-test model (**d**). *p<0.05, **p<0.01, ***p<0.001.

Supplemental Figure 2.

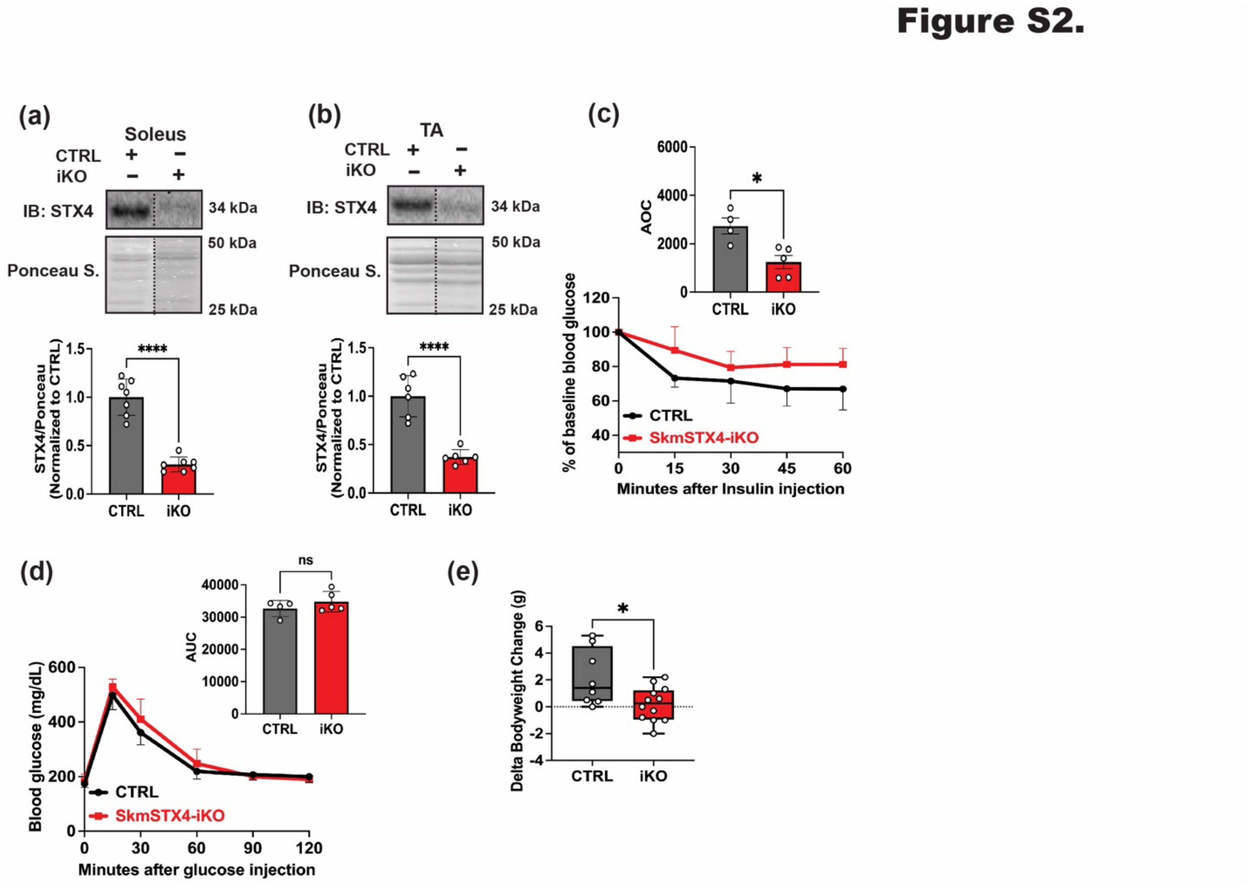


**Figure S2. Female skmSTX4-iKO mice present reduced insulin tolerance and bodyweight.**

Representative immunoblot (top) and quantification (bottom) of STX4 protein abundance across skeletal muscle depots (**a**) soleus and (**b**) Tibialis anterior from control (CTRL, grey) and skmSTX4-iKO (iKO, red) 20-week-old female mice normalized against total protein lysate stained with ponceau. (**c**) Intraperitoneal insulin tolerance test (IPITT) (bottom) of CTRL (black/grey) and skmSTX4-iKO (iKO, red) female 20-week-old mice with area over the curve (AOC) bar chart. (**d**) Intraperitoneal glucose tolerance test (IPGTT) between Ctrl (Black/Grey) and skmSTX4-iKO (Red) 20-week-old female mice with area under the curve (AUC) bar chart **(e)** Delta bodyweight change of initial weight to final weight of control (CTRL, grey) and skmSTX4-iKO (iKO, red) 20-week-old female mice. Data represents n = 4-12 mice per group in bar and line graphs shown as mean ± SD. Statistical significance was determined by unpaired two tailed Student's t test (**a, b, c** (AOC), **d** (AUC), **e**) and multiple t-test model (**c, d**). *p<0.05, ****p<0.0001, ns = not significant. Vertical dashed lines indicate the splicing of lanes within the same gel exposure. IB: Immunoblot, Ponceau S.: Ponceau stain.

Supplemental Figure 3.

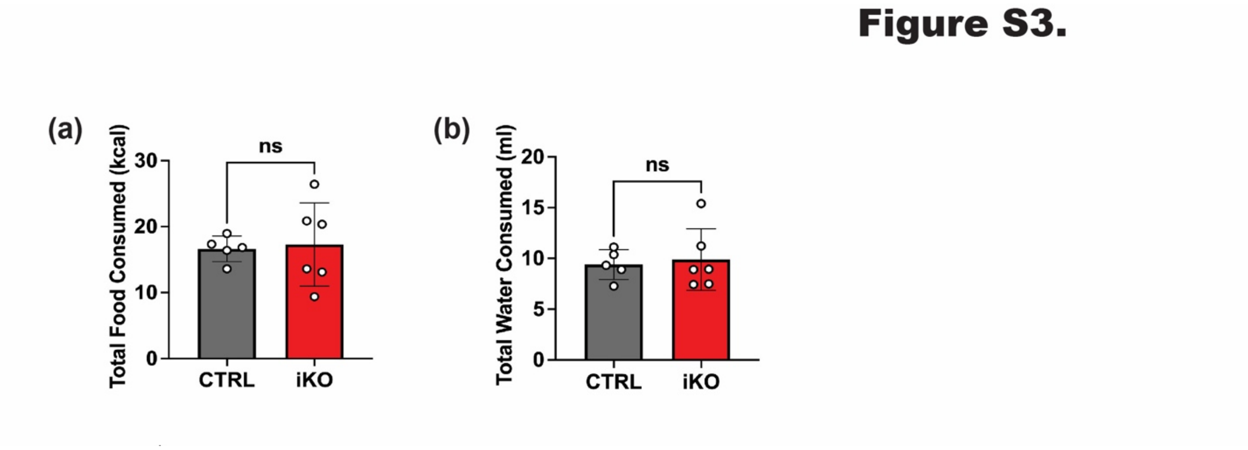


**Figure S3. STX4 ablation has no impact on food/water consumption in male skmSTX4-iKO mice.**

Metabolic caging analyses of the 26-week-old control (CTRL, black) and skmSTX4-iKO (iKO, red) male mice 72 hours period (n=5-6 mice/group) for (**a**) Total food consumed and (**b**) Total water consumed. Bar graph represents mean ± SD, unpaired two-tailed Student's *t*-test, ns = not significant.

Supplemental Figure 4.

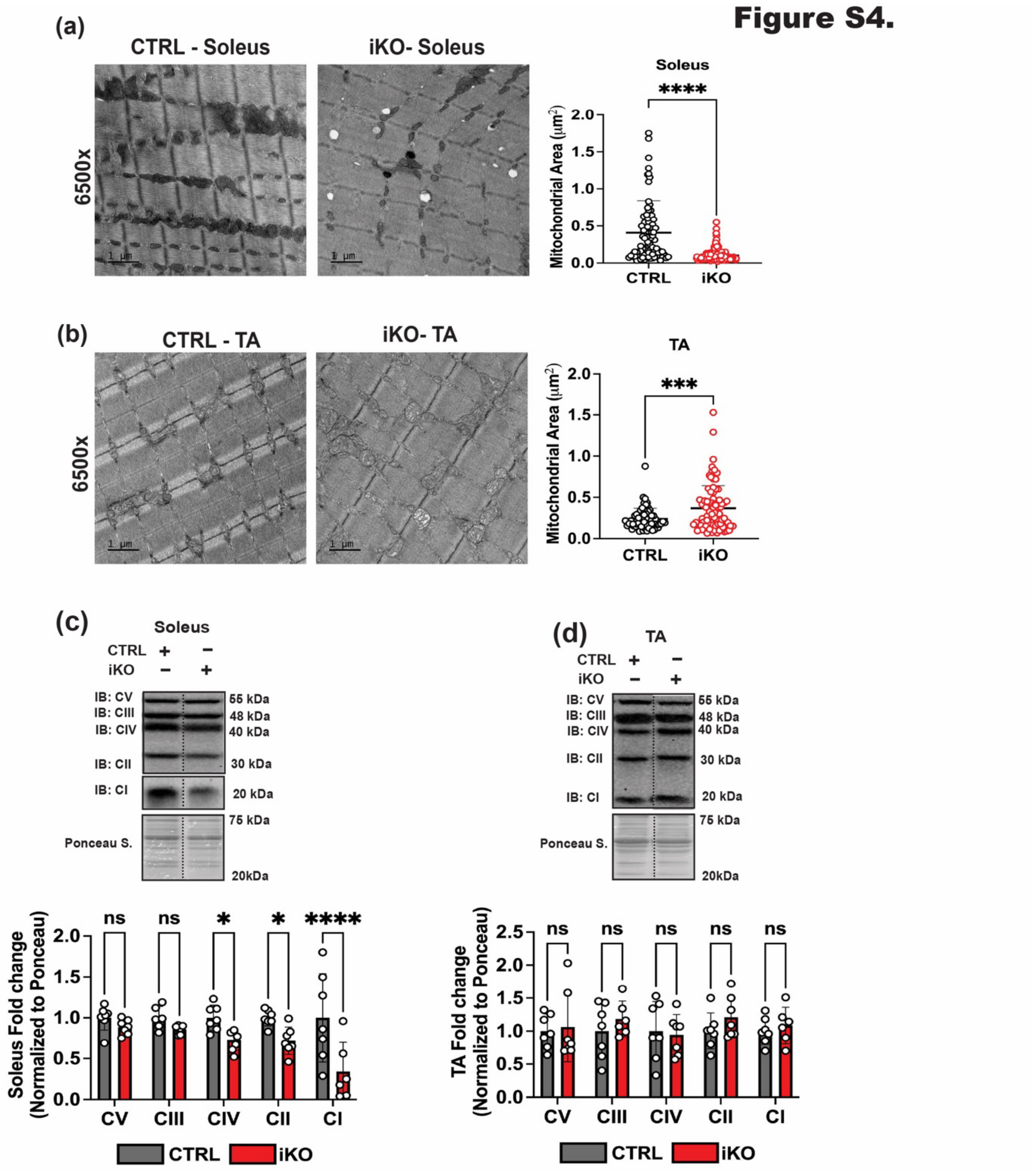


**Figure S4. STX4 ablation has a heterogenous impact on mitochondrial structure and electron transport chain complex abundance between oxidative rich soleus and glycolytic rich TA muscle in male skmSTX4-iKO mice.** Representative transmission electron microscopy images and mitochondrial area (μm$^2$) quantification of mitochondrial from **(a)** Soleus (n = 96-202 total mitochondria) and **(b)** TA muscle (n = 63-85 total mitochondria) from 20-week-old control (CTRL, Grey) and skmSTX4-iKO (iKO, Red) female mice. Black bar = 1 μm (6500X).

### Supplemental Figure 4.

Representative immunoblot (top) and quantification (bottom) of total electron transport chain (ETC) complexes V (CV), III (CIII), IV (CIV), II (CII) and I (CI) normalized against total protein lysate stained with ponceau in **(c)** soleus and **(d)** TA muscle from 20-week-old control (CTRL, Grey) and skmSTX4-iKO (iKO, Red) female mice (n=3-6 mice/group). Data in bar graphs shown as mean ± SD. Statistical significance was determined by unpaired two tailed Student's *t* test (**a, b**) and two-way ANOVA with uncorrected Fisher's LSD test (**c, d**). *p<0.05, ***p<0.001, ****p<0.0001, ns = not significant. Vertical dashed lines indicate the splicing of lanes within the same gel exposure. IB: Immunoblot, Ponceau S.: Ponceau stain.

Supplemental Figure 5.

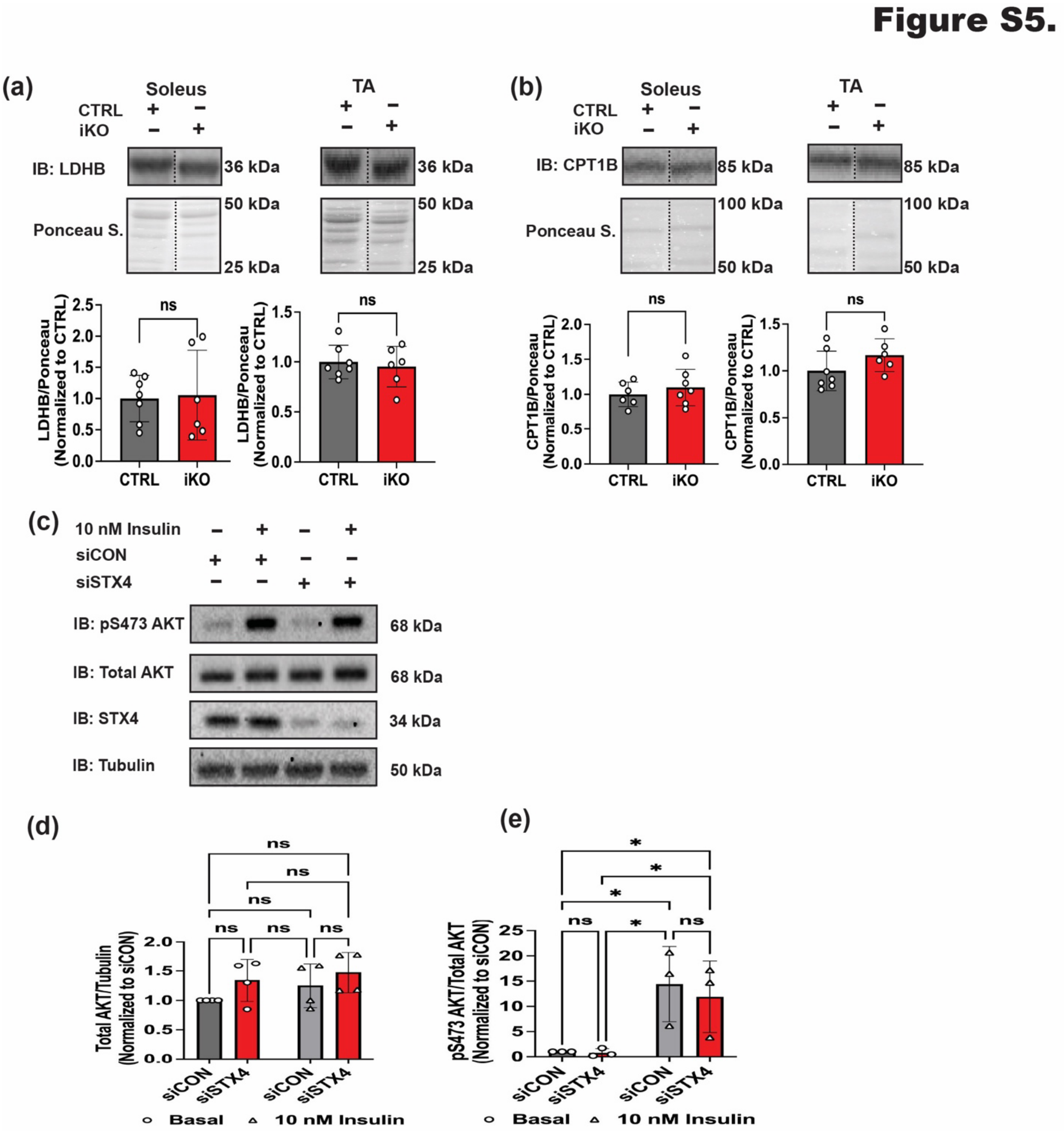


**Figure S5. STX4 ablation does not impact markers for lactate conversion, fatty acid oxidation or canonical insulin signaling in skeletal muscle.** Representative immunoblot (top) and quantification (bottom) of (**a**) LDHB and (**b**) CPT1B normalized against total protein lysate with ponceau in soleus and Tibialis anterior (TA) muscle from control (CTRL, Grey) and skmSTX4-iKO (iKO, Red) 20-week-old male mice (n=7 mice/group). (**c**) Representative immunoblot of pS473 AKT, total AKT, STX4 and Tubulin with quantification for (**d**) total AKT and (**e**) pS473 AKT normalized against Tubulin in siCON (Grey) and siSTX4 (Red) L6.GLUT4myc myotubes with or without 10 nM insulin stimulation (n=3-4 independent passages). Data in bar graphs shown as mean ± SD. Statistical significance was determined by unpaired two tailed Student's *t* test (**a, b**) and two-way ANOVA with uncorrected Fisher's LSD test (**c, d**). *$p<0.05$, ns = not significant. Vertical dashed lines indicate the splicing of lanes within the same gel exposure. IB: Immunoblot, Ponceau S.: Ponceau stain.

Supplemental Figure 6.

Figure S6.

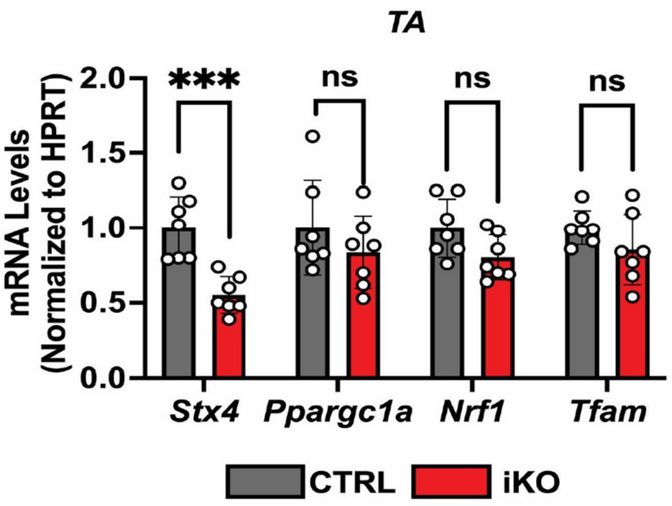


**Figure S6. STX4 ablation has no impact on mitochondrial biogenesis genes in the TA muscle in male skmSTX4-iKO mice.** Quantification of mRNA levels via qPCR of *Stx4*, *Ppargc1a* (Pgc1-α), *Nrf1* and *Tfam* normalized against housekeeping gene *Hprt* in the Tibialis anterior (TA) muscle of 20-week-old CTRL (Grey) and skmSTX4-iKO (Red) male mice. Data represents n = 7 mice per group in bar graphs shown as mean ± SD. Statistical significance was determined two-way ANOVA with uncorrected Fisher's LSD test, ***p<0.001, ns = not significant.
