## Supplementary material for "STX4 is indispensable for mitochondrial homeostasis in skeletal muscle": Raw Blots

### Raw Blots File

**Fig. 1a**

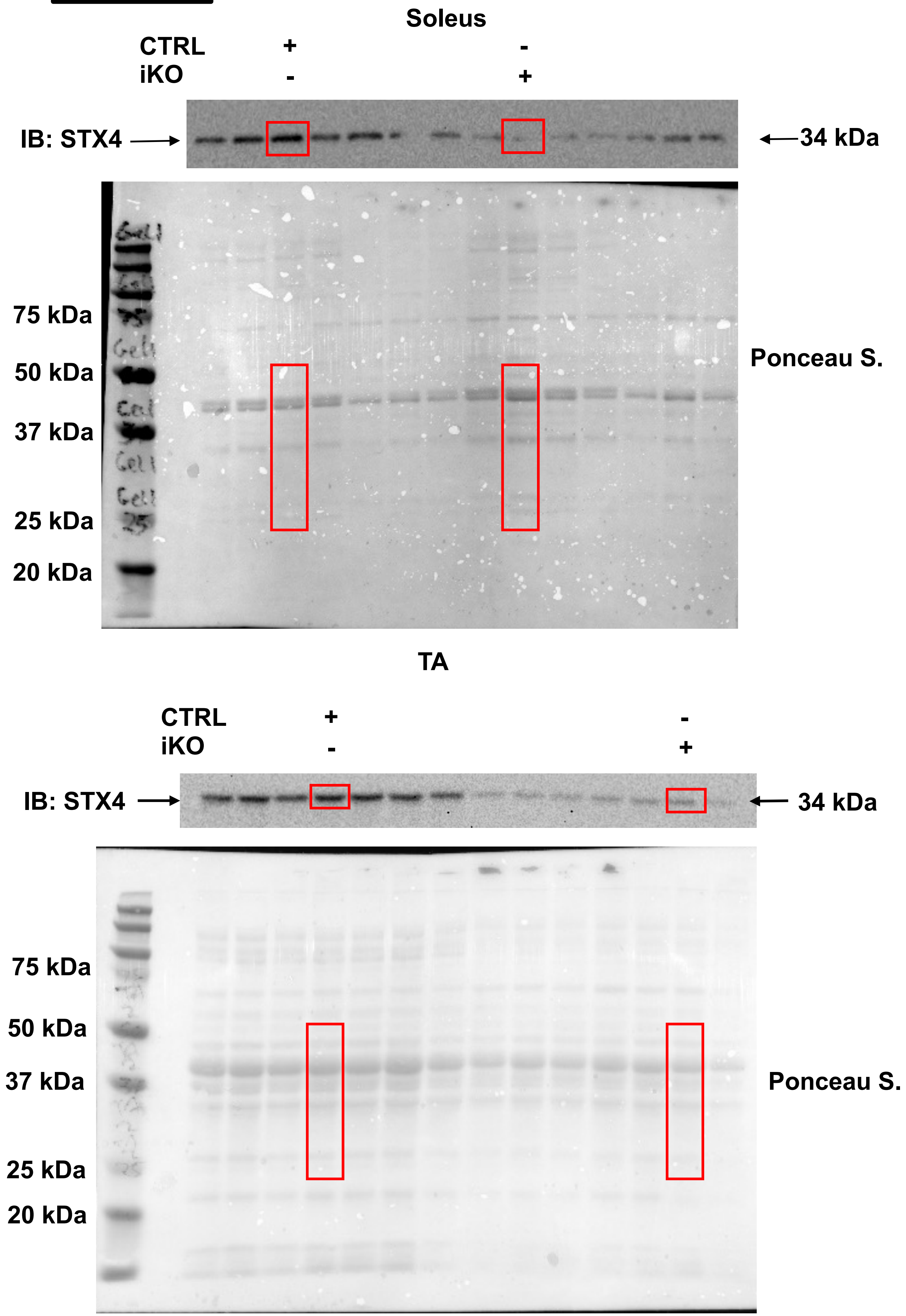
Soleus
CTRL + -
iKO - +
IB: STX4
34 kDa
75 kDa
50 kDa
37 kDa
25 kDa
20 kDa
Ponceau S.
TA
CTRL + -
iKO - +
IB: STX4
34 kDa
75 kDa
50 kDa
37 kDa
25 kDa
20 kDa
Ponceau S.

**Fig. 1a**

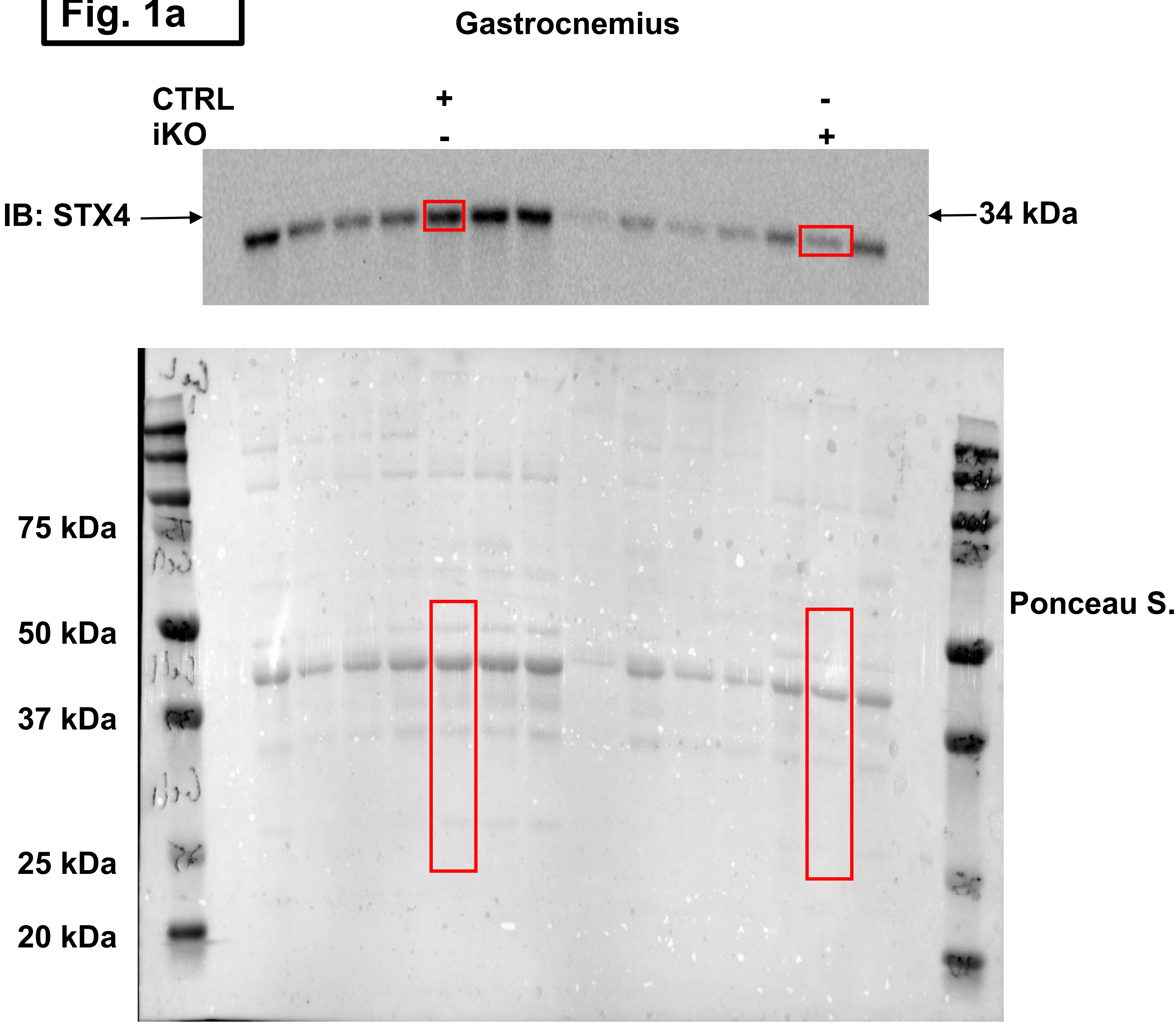
Gastrocnemius
CTRL + -
iKO - +
IB: STX4
34 kDa
75 kDa
50 kDa
37 kDa
25 kDa
20 kDa
Ponceau S.

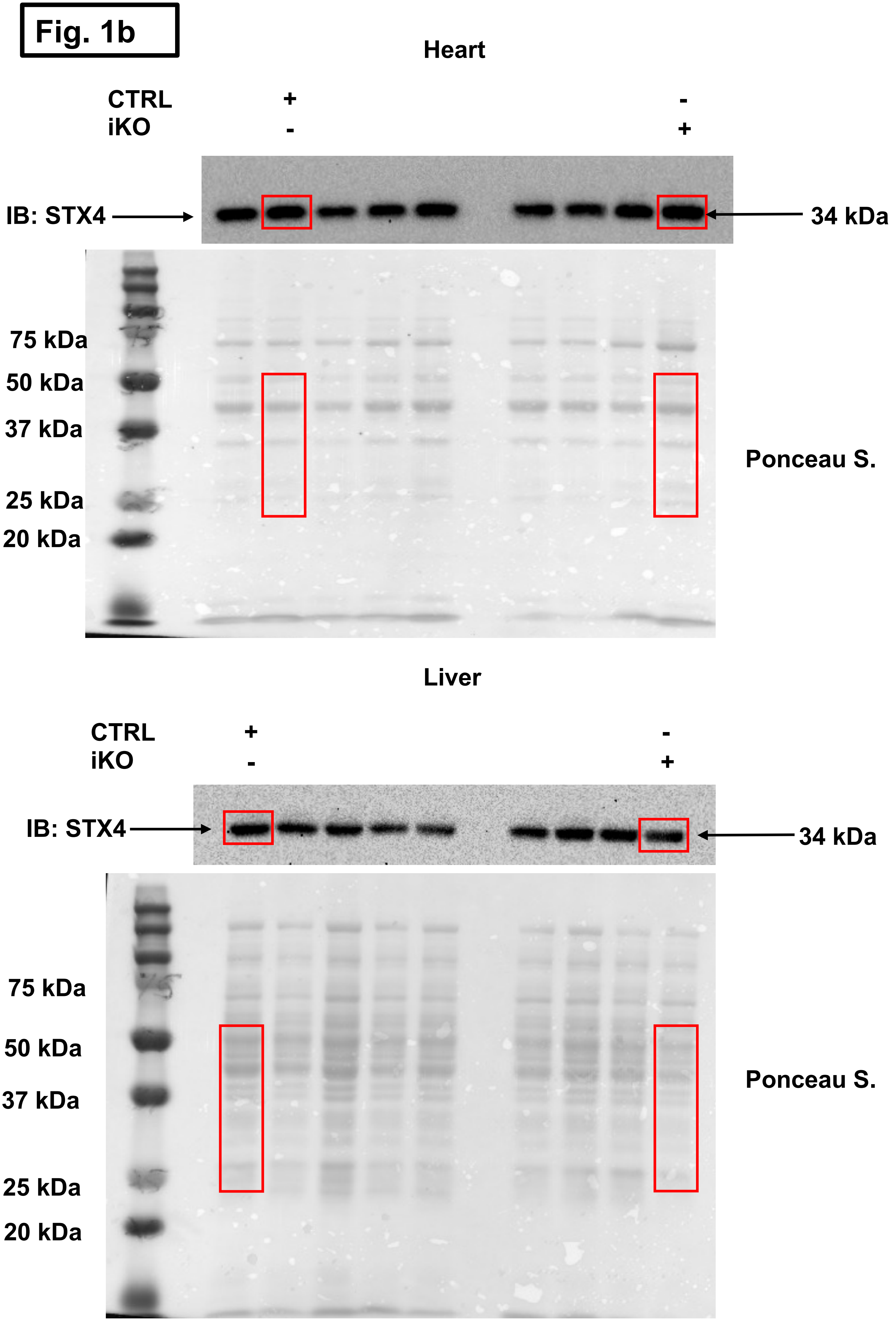
Fig. 1b
Heart
CTRL + -
iKO - +
IB: STX4
34 kDa
75 kDa
50 kDa
37 kDa
25 kDa
20 kDa
Ponceau S.
Liver
CTRL + -
iKO - +
IB: STX4
34 kDa
75 kDa
50 kDa
37 kDa
25 kDa
20 kDa
Ponceau S.

**Fig. 3c**

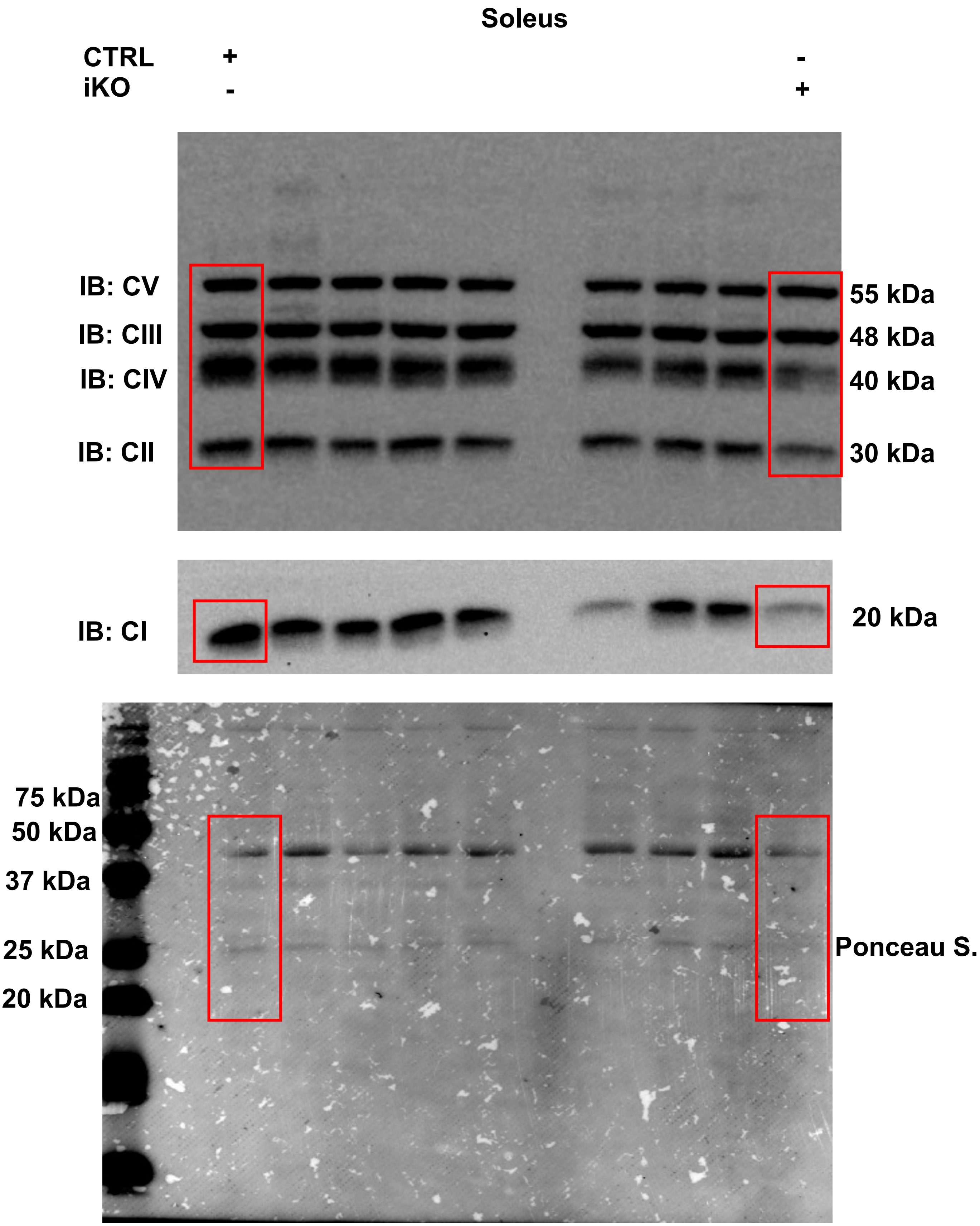
Soleus
CTRL + -
iKO - +
IB: CV
IB: CIII
IB: CIV
IB: CII
55 kDa
48 kDa
40 kDa
30 kDa
IB: CI
20 kDa
75 kDa
50 kDa
37 kDa
25 kDa
20 kDa
Ponceau S.

Fig. 3d

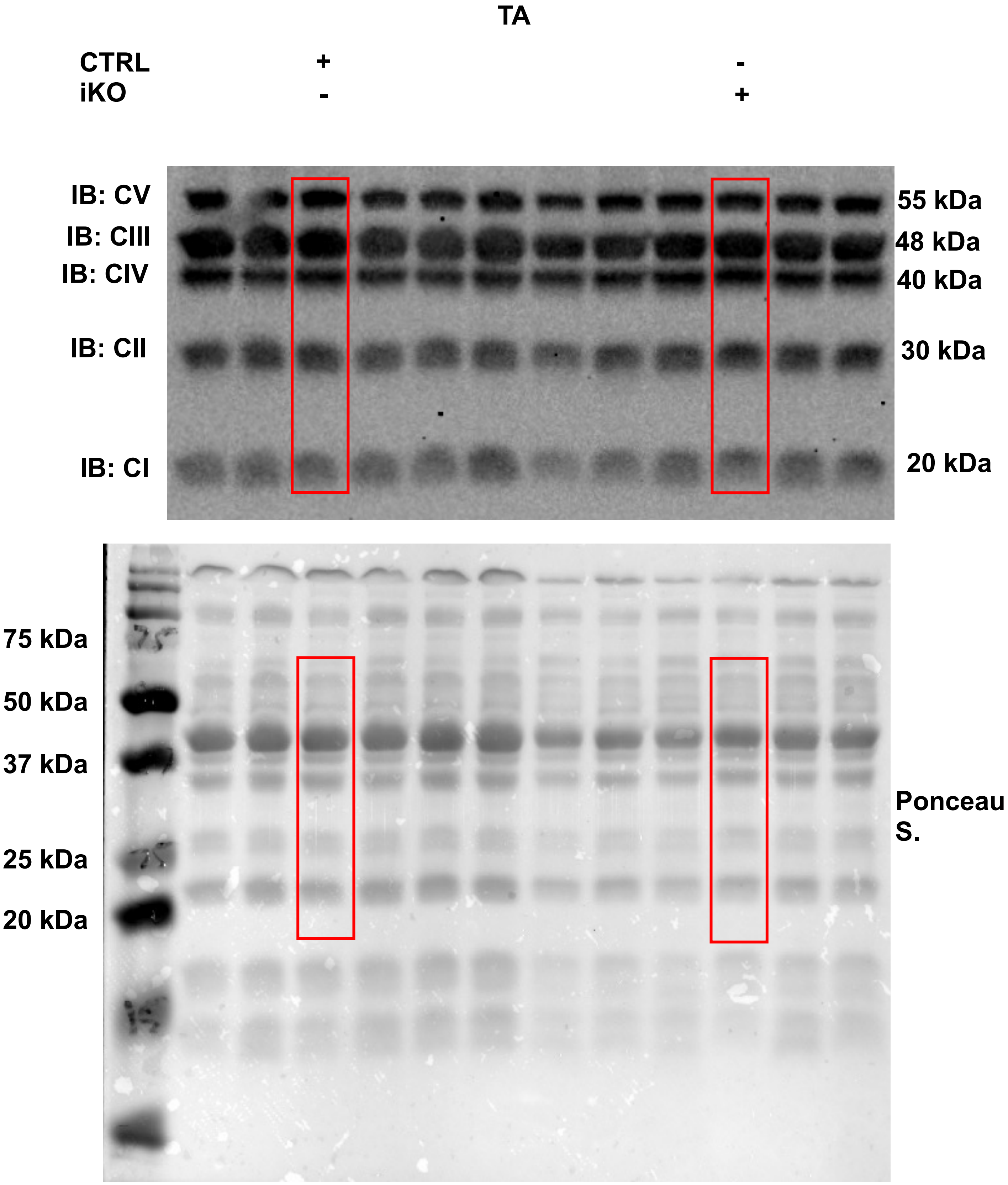
TA
CTRL + -
iKO - +
IB: CV
IB: CIII
IB: CIV
IB: CII
IB: CI
55 kDa
48 kDa
40 kDa
30 kDa
20 kDa
75 kDa
50 kDa
37 kDa
25 kDa
20 kDa
Ponceau S.

Fig. 4b

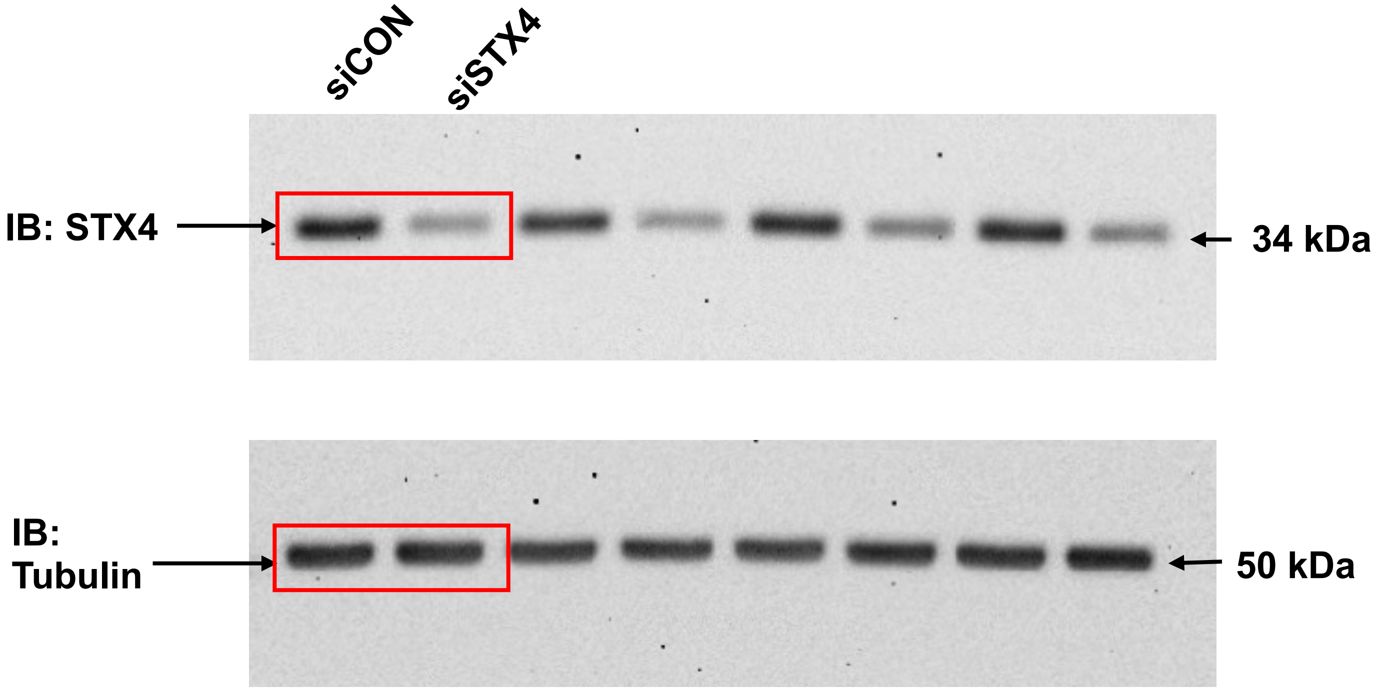

Fig. 4c

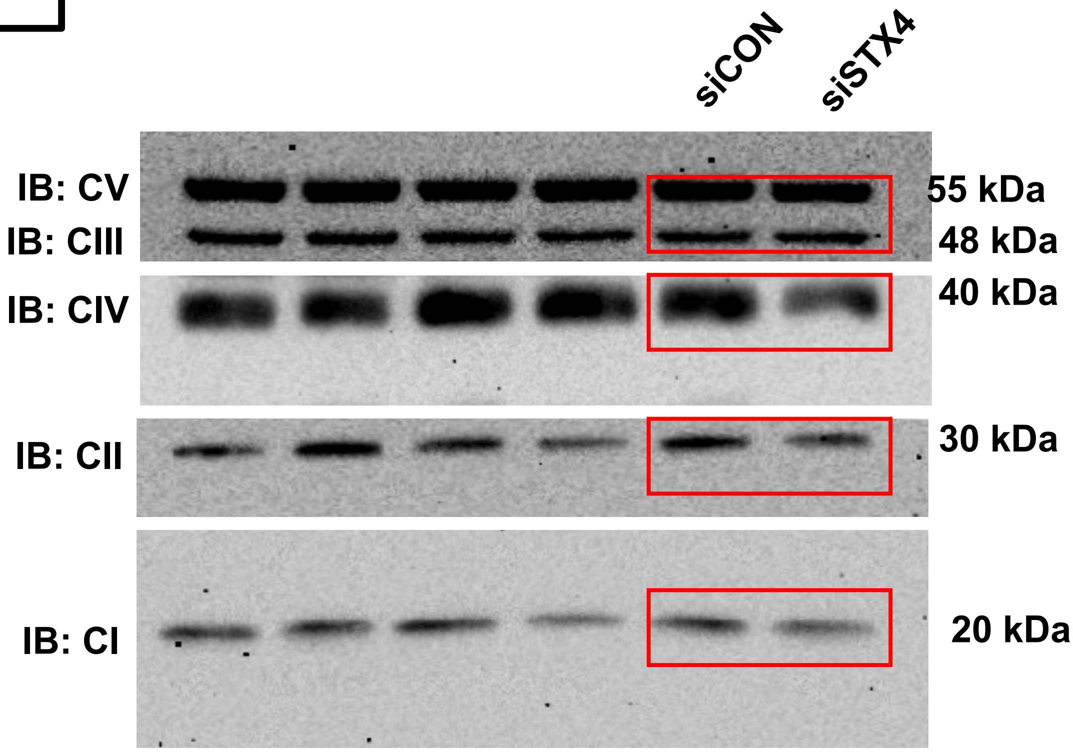
siCON
siSTX4
IB: CV
IB: CIII
IB: CIV
IB: CII
IB: CI
55 kDa
48 kDa
40 kDa
30 kDa
20 kDa

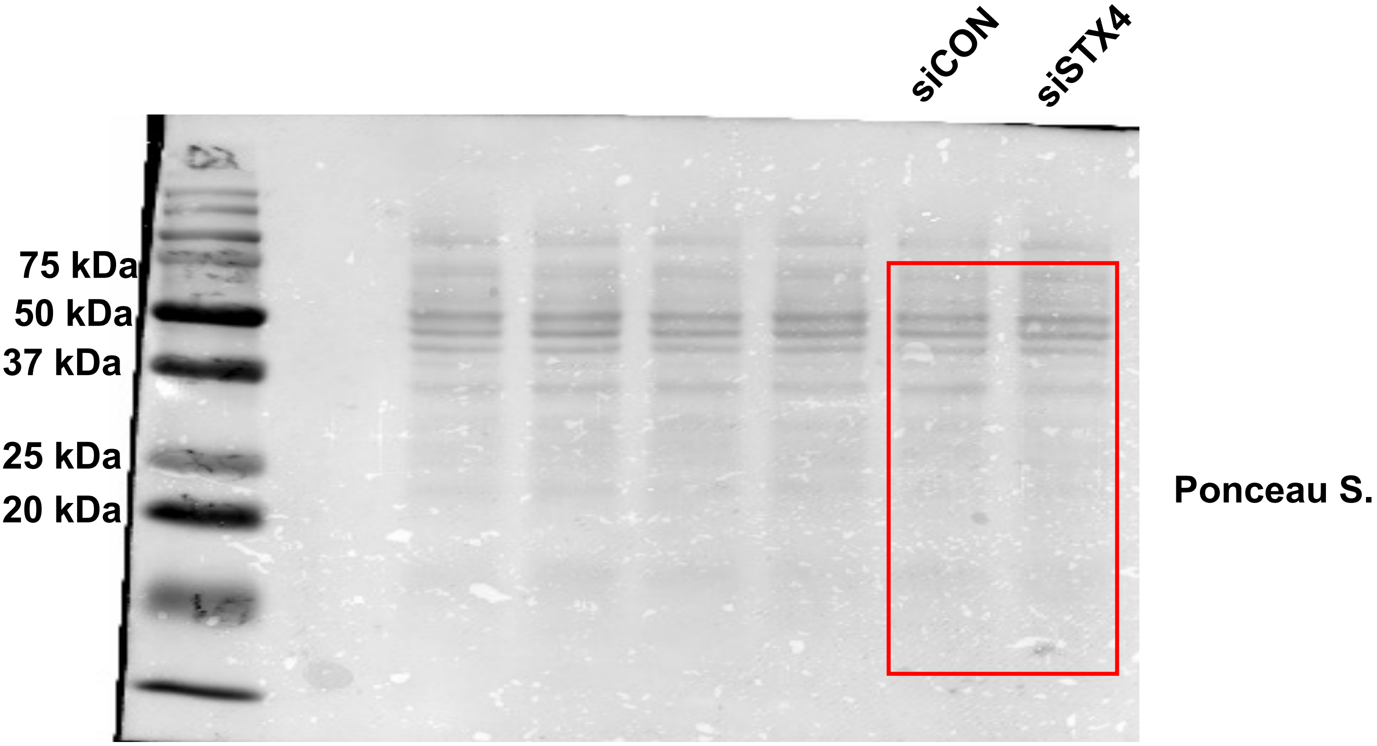
siCON
siSTX4
75 kDa
50 kDa
37 kDa
25 kDa
20 kDa
Ponceau S.

**Fig. 5e**

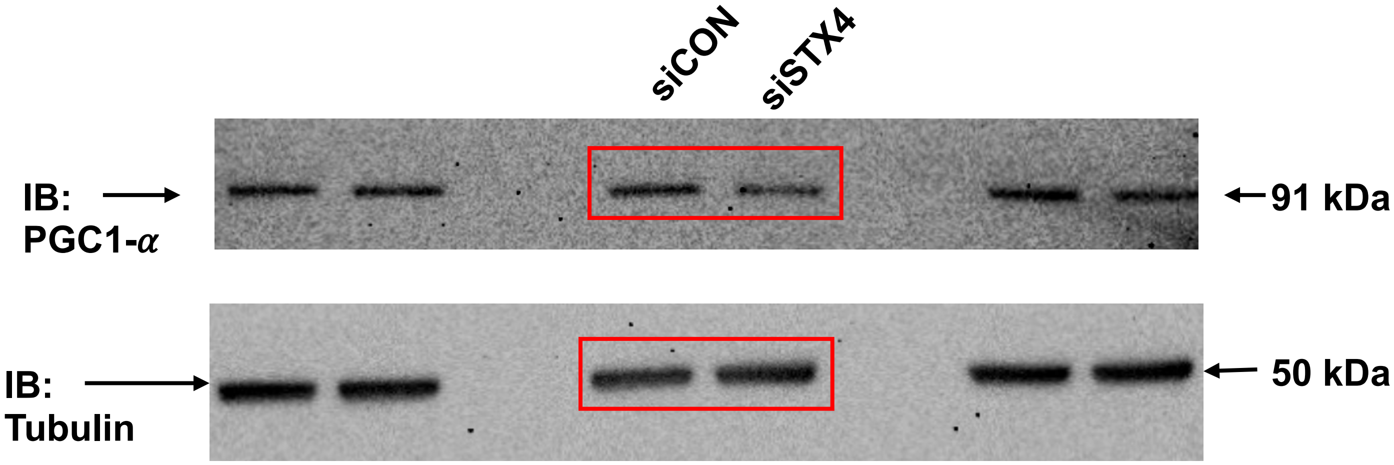

**Fig. 5f**

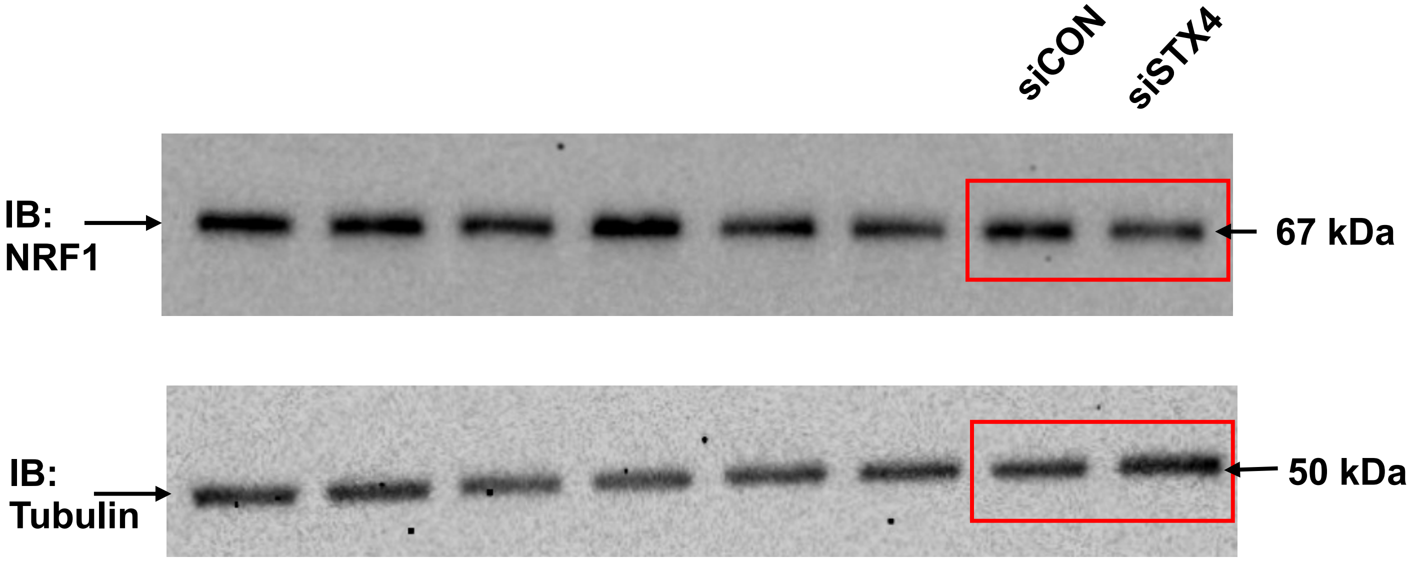

#### Fig. 7a

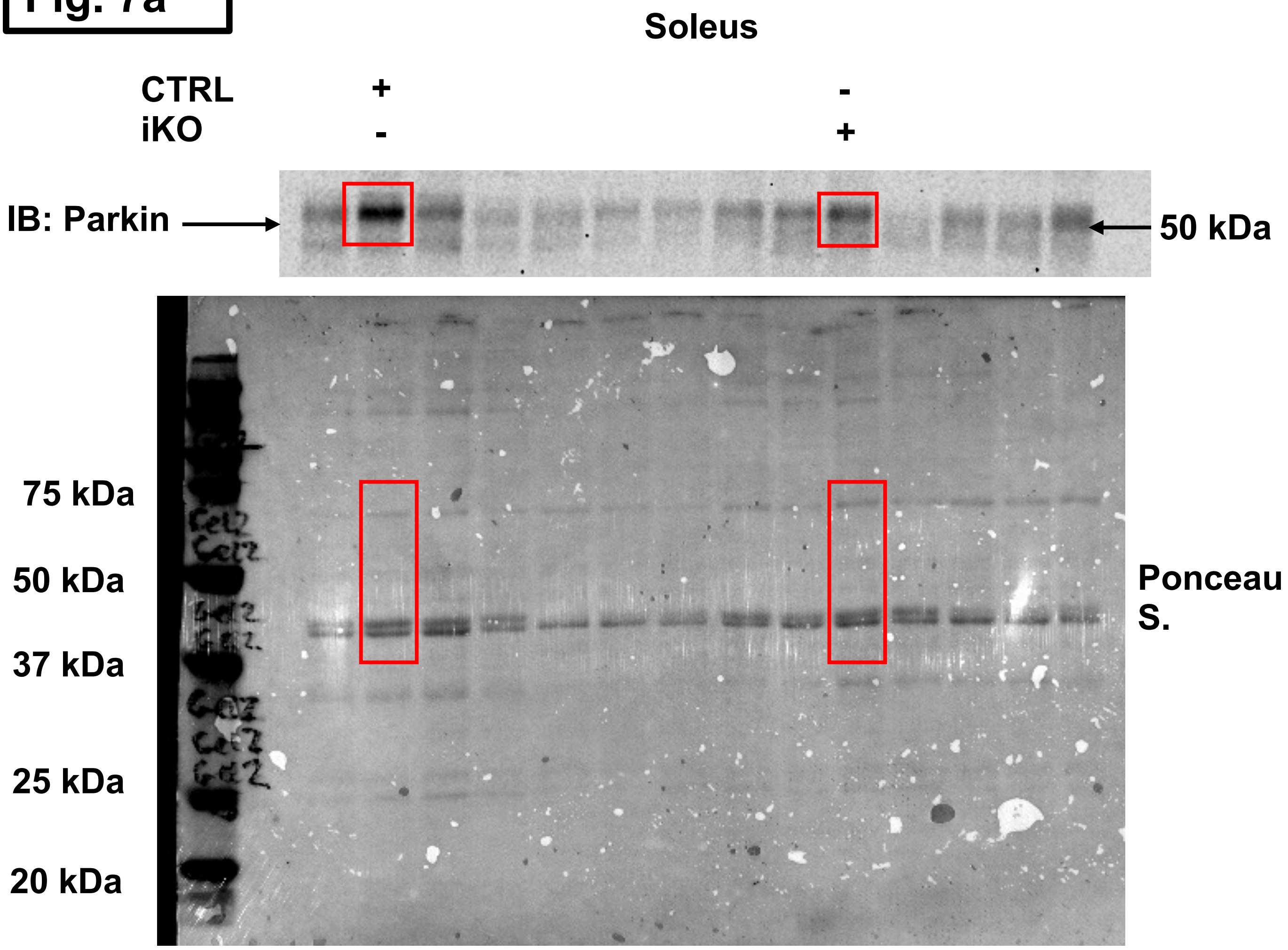

Soleus
CTRL + -
iKO - +
IB: Parkin
50 kDa
75 kDa
50 kDa
37 kDa
25 kDa
20 kDa
Ponceau S.

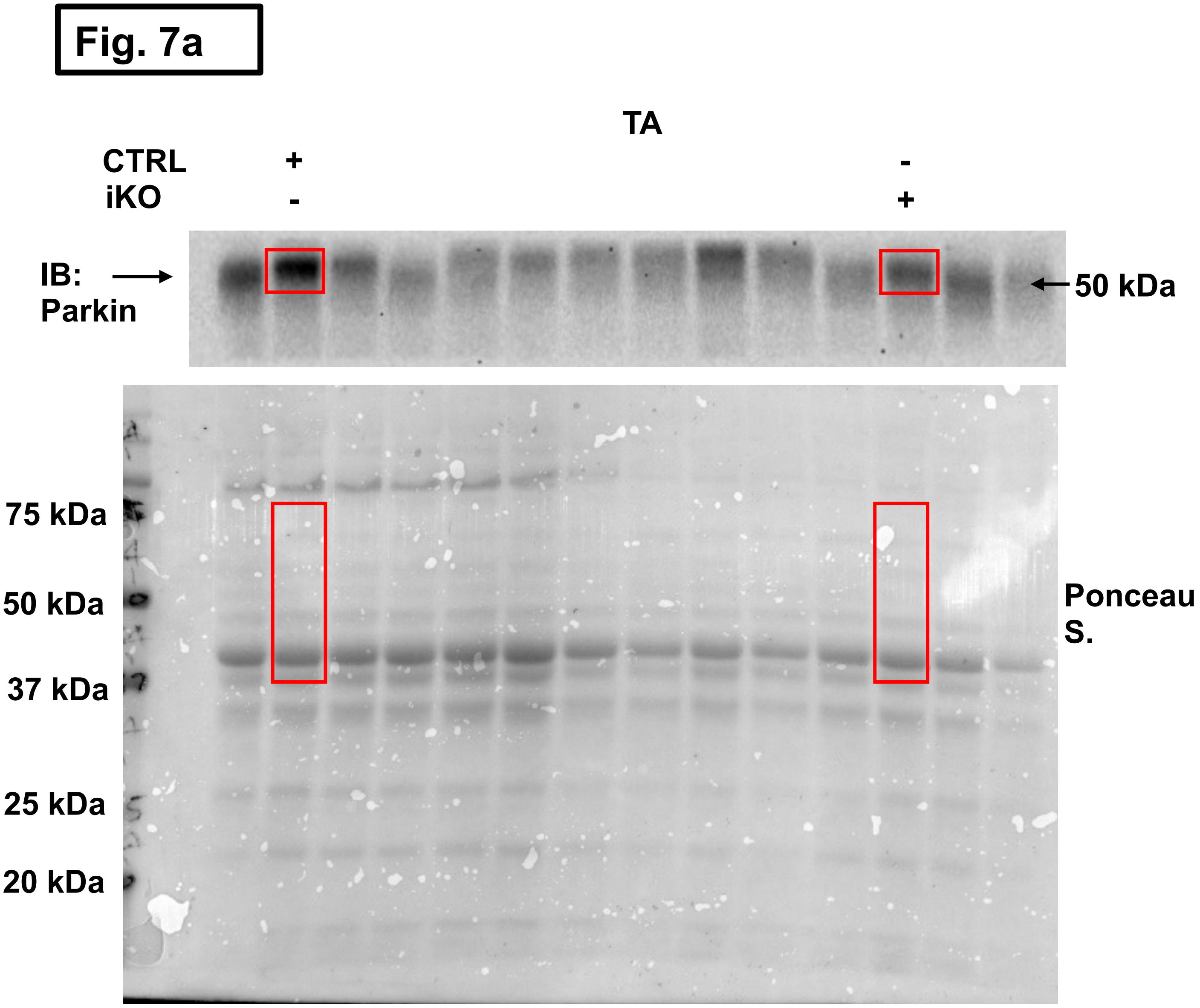

Fig. 7a
TA
CTRL + -
iKO - +
IB: Parkin
50 kDa
75 kDa
50 kDa
37 kDa
25 kDa
20 kDa
Ponceau S.

Fig. 7b

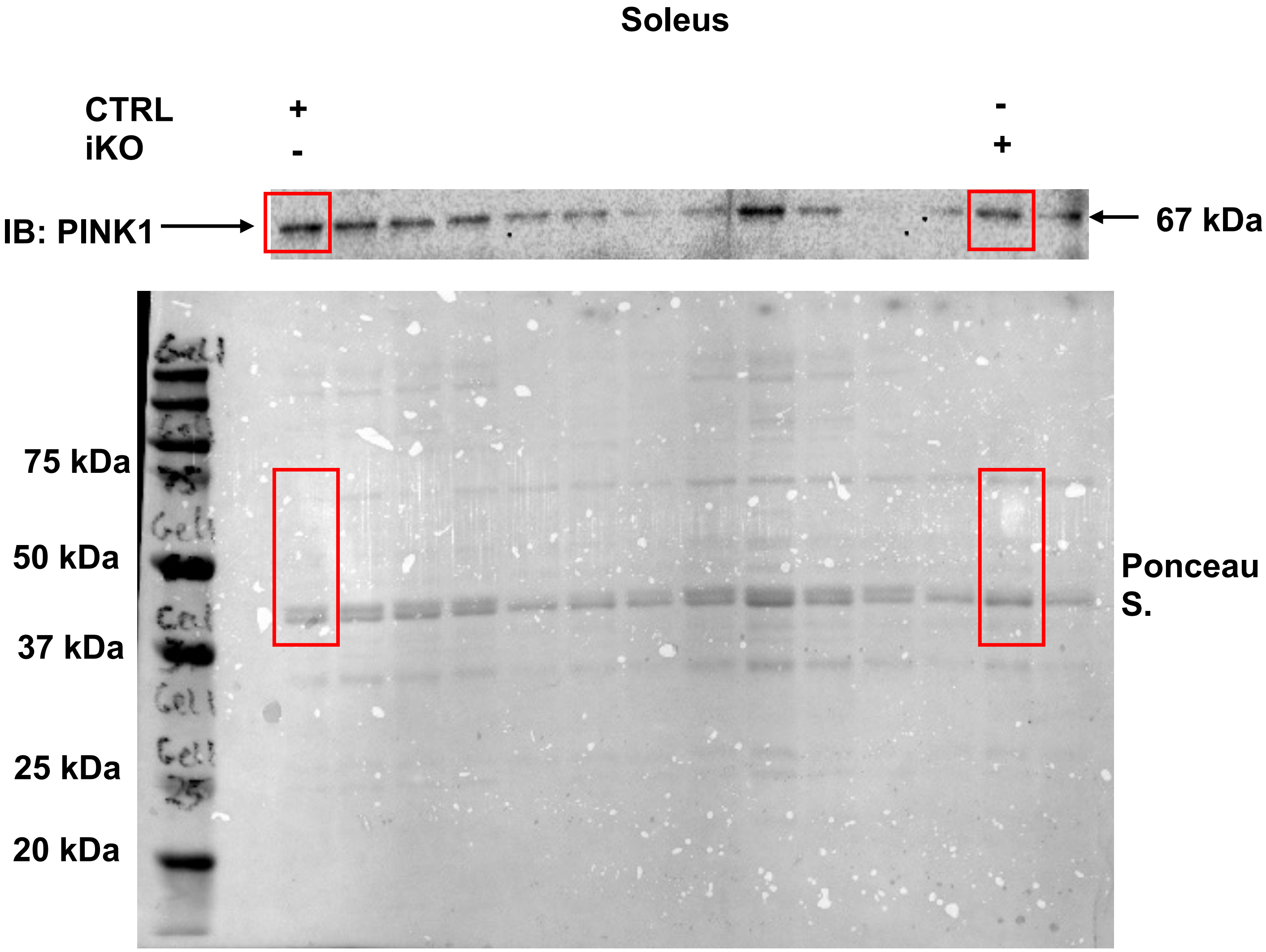
Soleus
CTRL
iKO
+
-
-
+
IB: PINK1
67 kDa
75 kDa
50 kDa
37 kDa
25 kDa
20 kDa
Ponceau S.

**Fig. 7b**

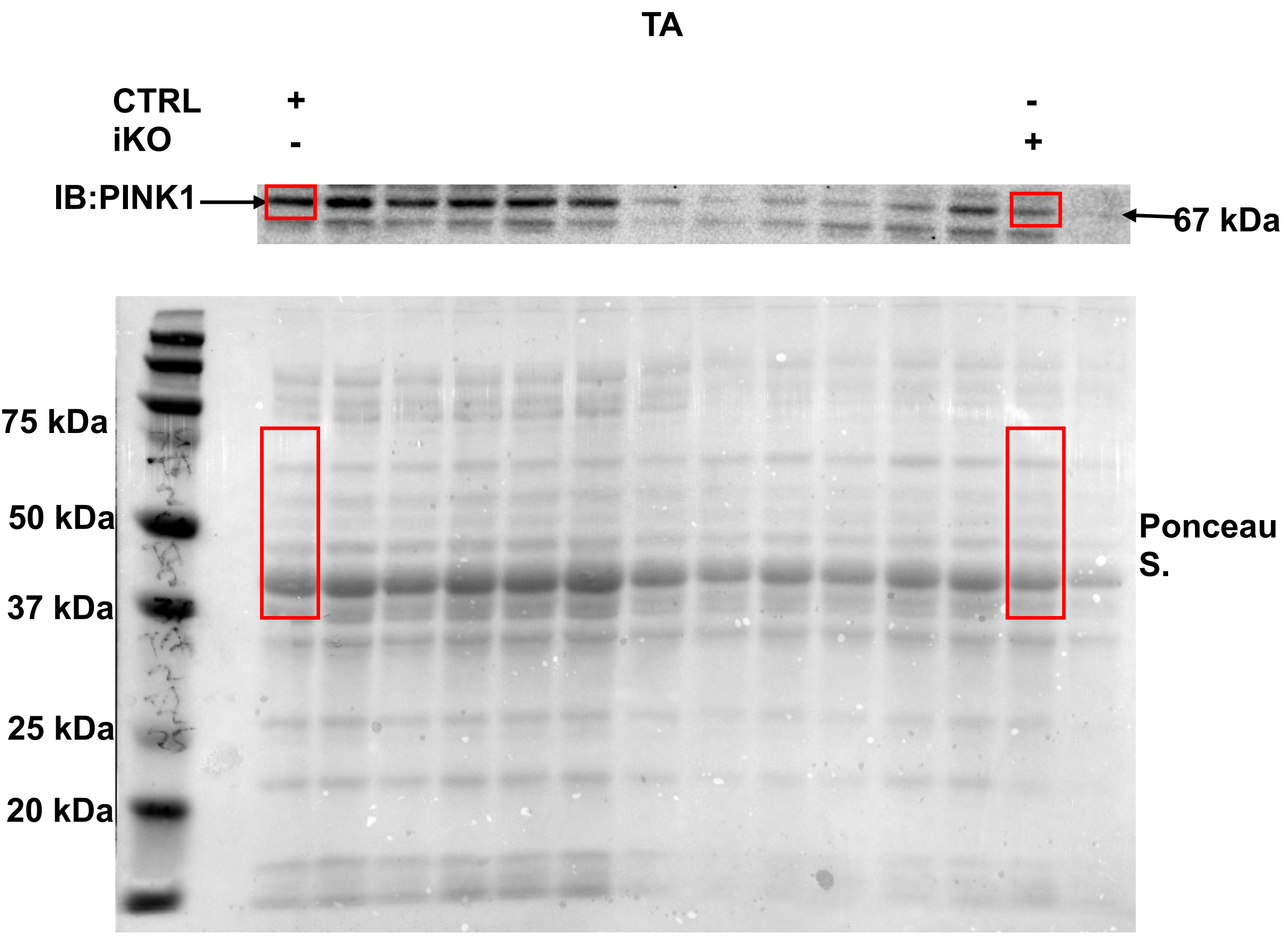
TA
CTRL
iKO
+
-
-
+
IB:PINK1
67 kDa
75 kDa
50 kDa
37 kDa
25 kDa
20 kDa
Ponceau S.

Fig. 7c

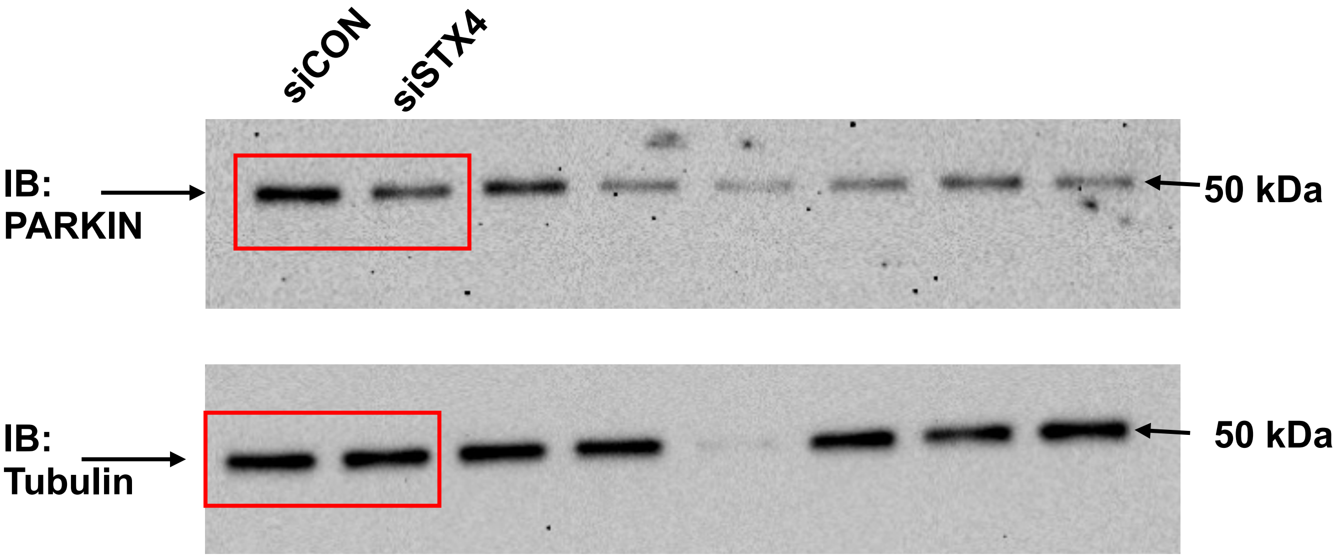

Fig. 7d

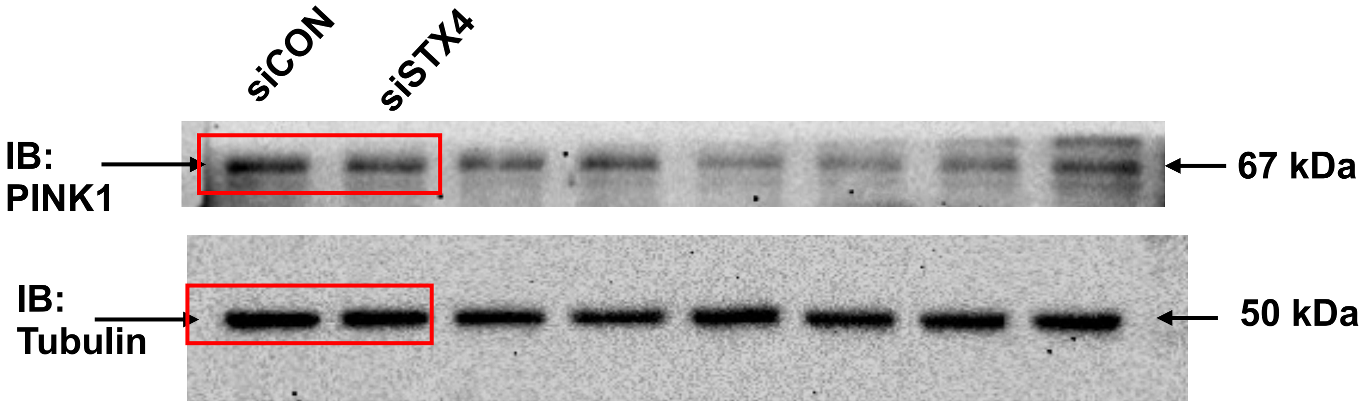

**Fig. S2a**

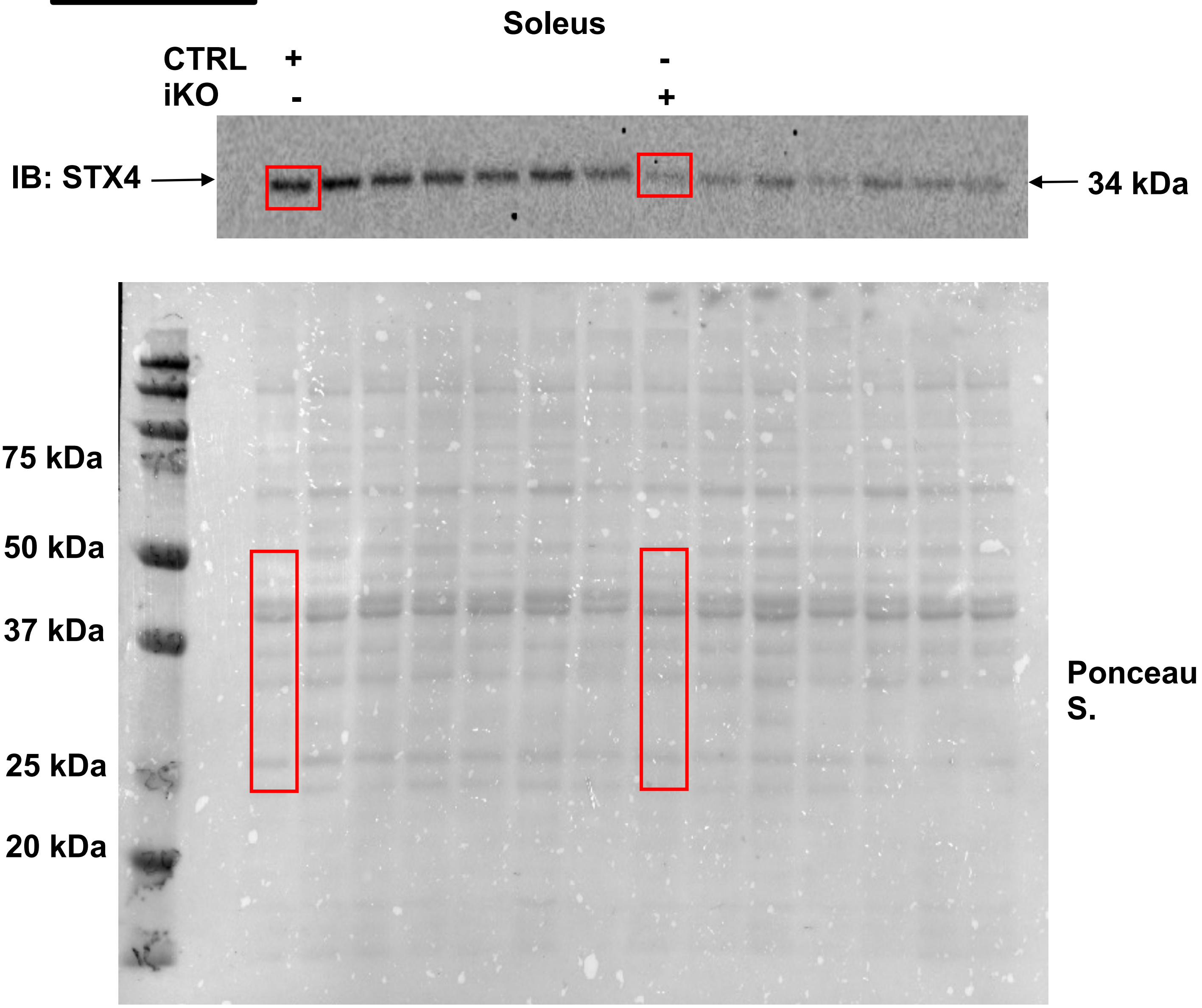

Soleus
CTRL + -
iKO - +
IB: STX4
34 kDa
75 kDa
50 kDa
37 kDa
25 kDa
20 kDa
Ponceau S.

**Fig. S2b**

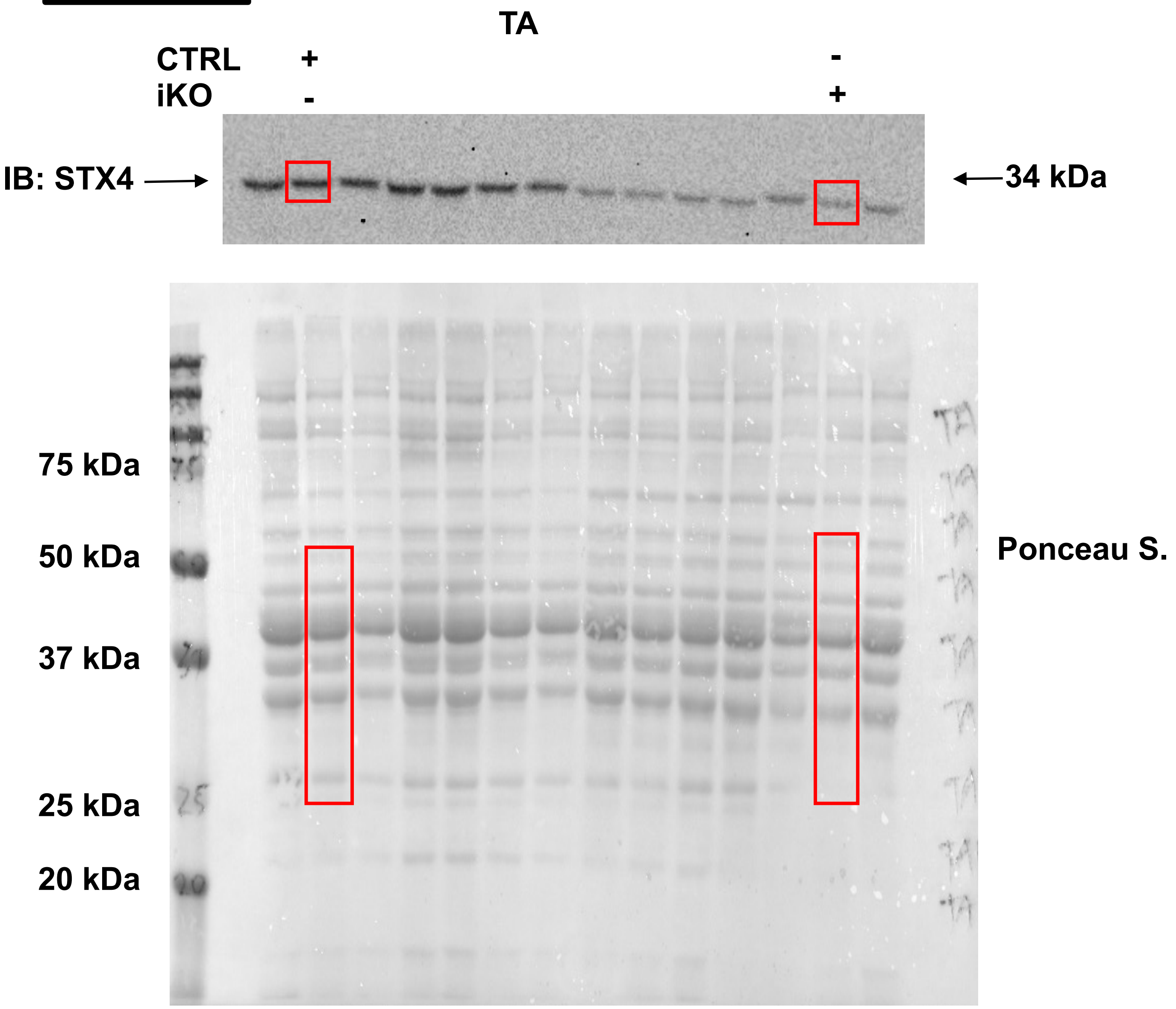
TA
CTRL + -
iKO - +
IB: STX4
34 kDa
75 kDa
50 kDa
37 kDa
25 kDa
20 kDa
Ponceau S.

Fig. S4c

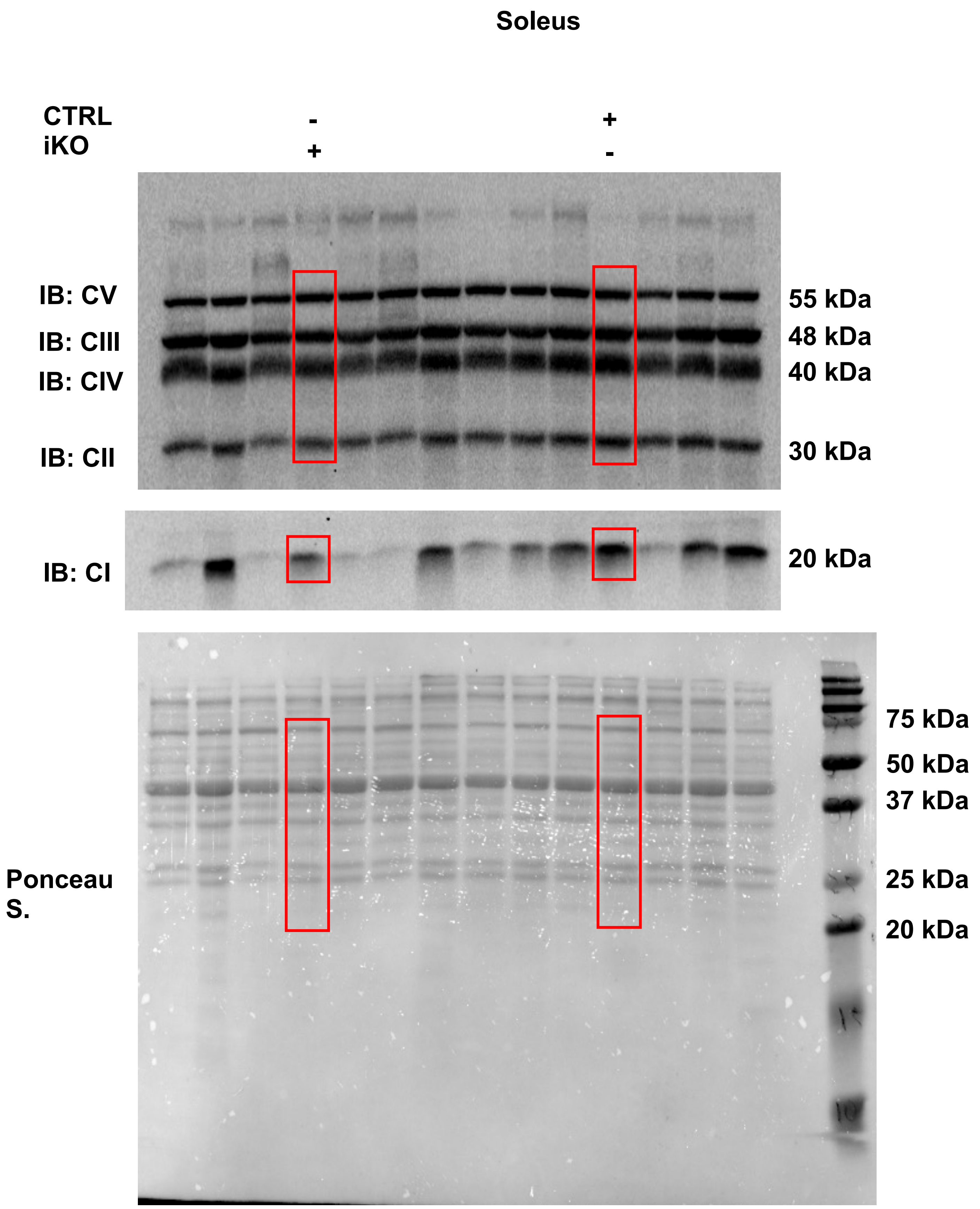
Soleus
CTRL
iKO
-
+
+
-
IB: CV
IB: CIII
IB: CIV
IB: CII
55 kDa
48 kDa
40 kDa
30 kDa
IB: CI
20 kDa
Ponceau S.
75 kDa
50 kDa
37 kDa
25 kDa
20 kDa

**Fig. S4d**

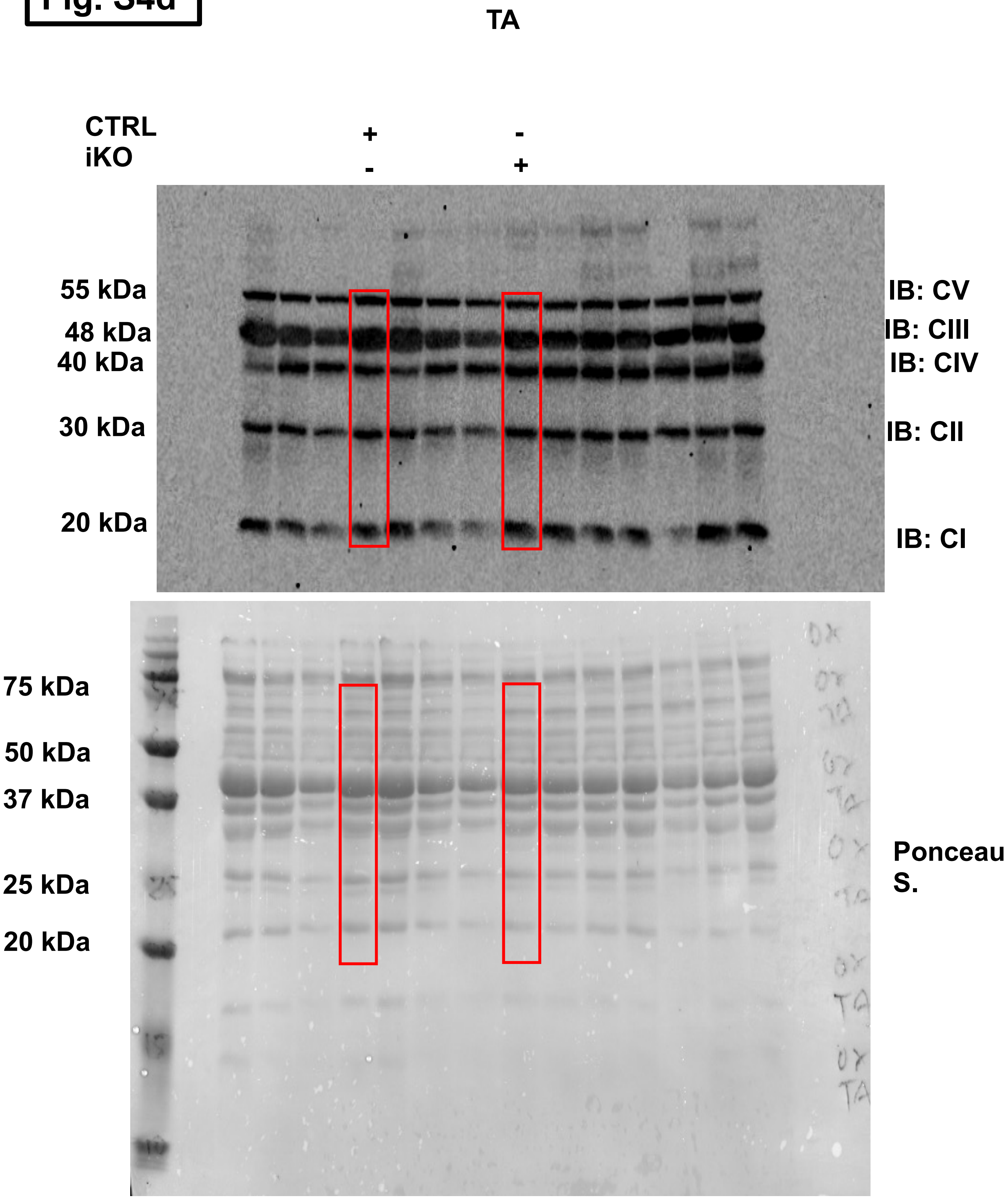
TA
CTRL + -
iKO - +
55 kDa
48 kDa
40 kDa
30 kDa
20 kDa
IB: CV
IB: CIII
IB: CIV
IB: CII
IB: CI
75 kDa
50 kDa
37 kDa
25 kDa
20 kDa
Ponceau S.

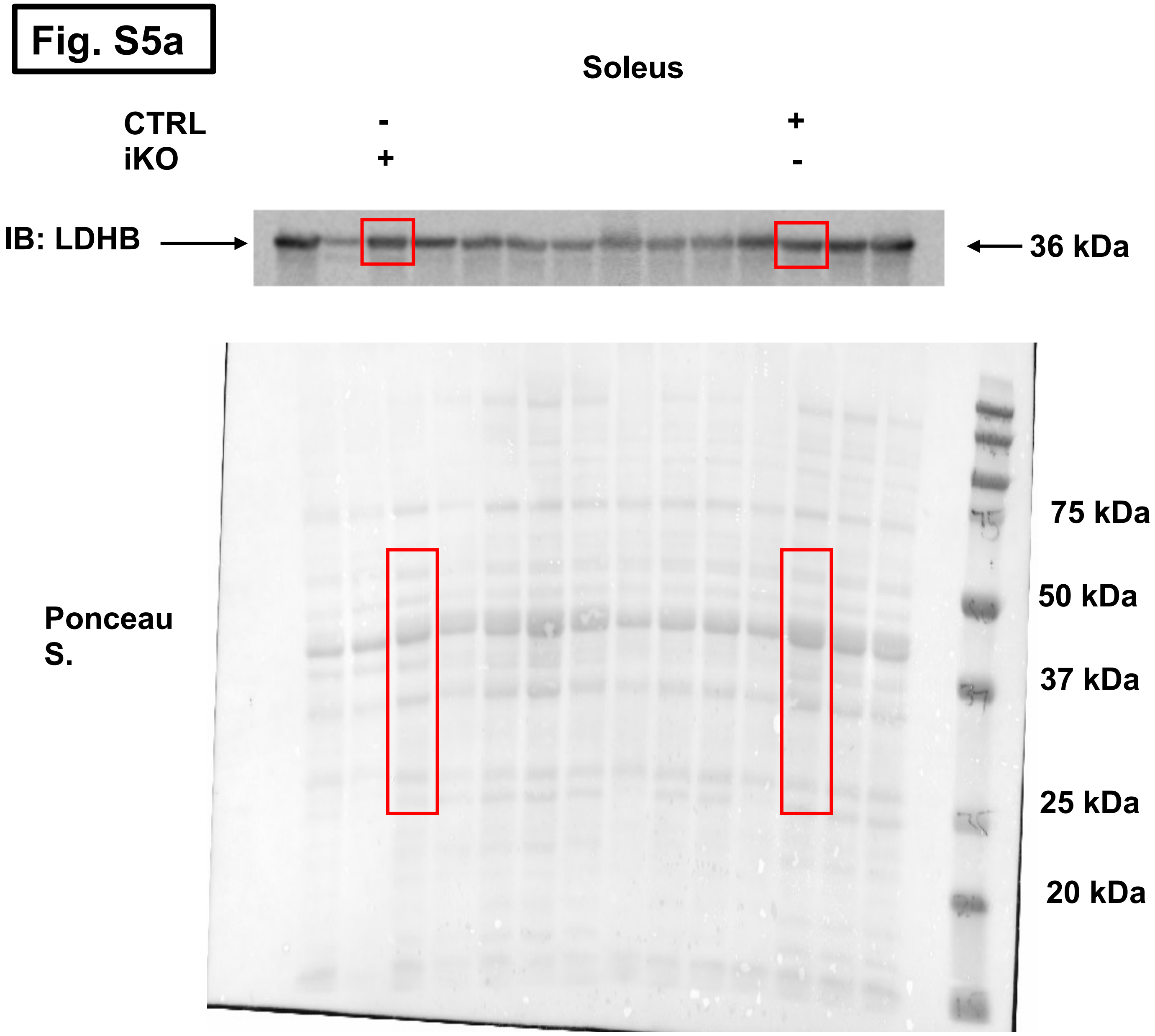
Fig. S5a
Soleus
CTRL
iKO
-
+
+
-
IB: LDHB
36 kDa
Ponceau S.
75 kDa
50 kDa
37 kDa
25 kDa
20 kDa

Fig. S5a

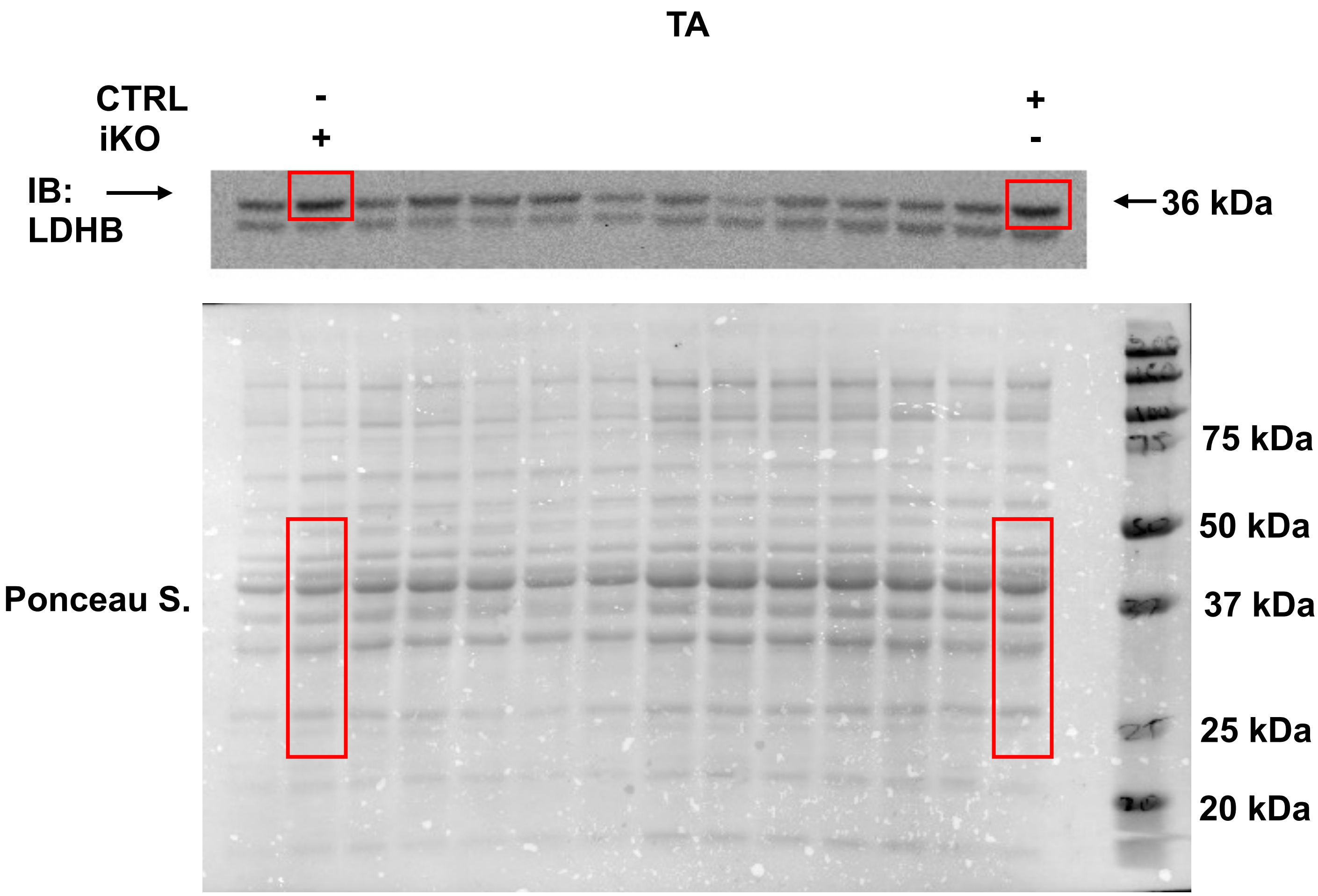
TA
CTRL
iKO
-
+
+
-
IB:
LDHB
36 kDa
Ponceau S.
75 kDa
50 kDa
37 kDa
25 kDa
20 kDa

Fig. S5b

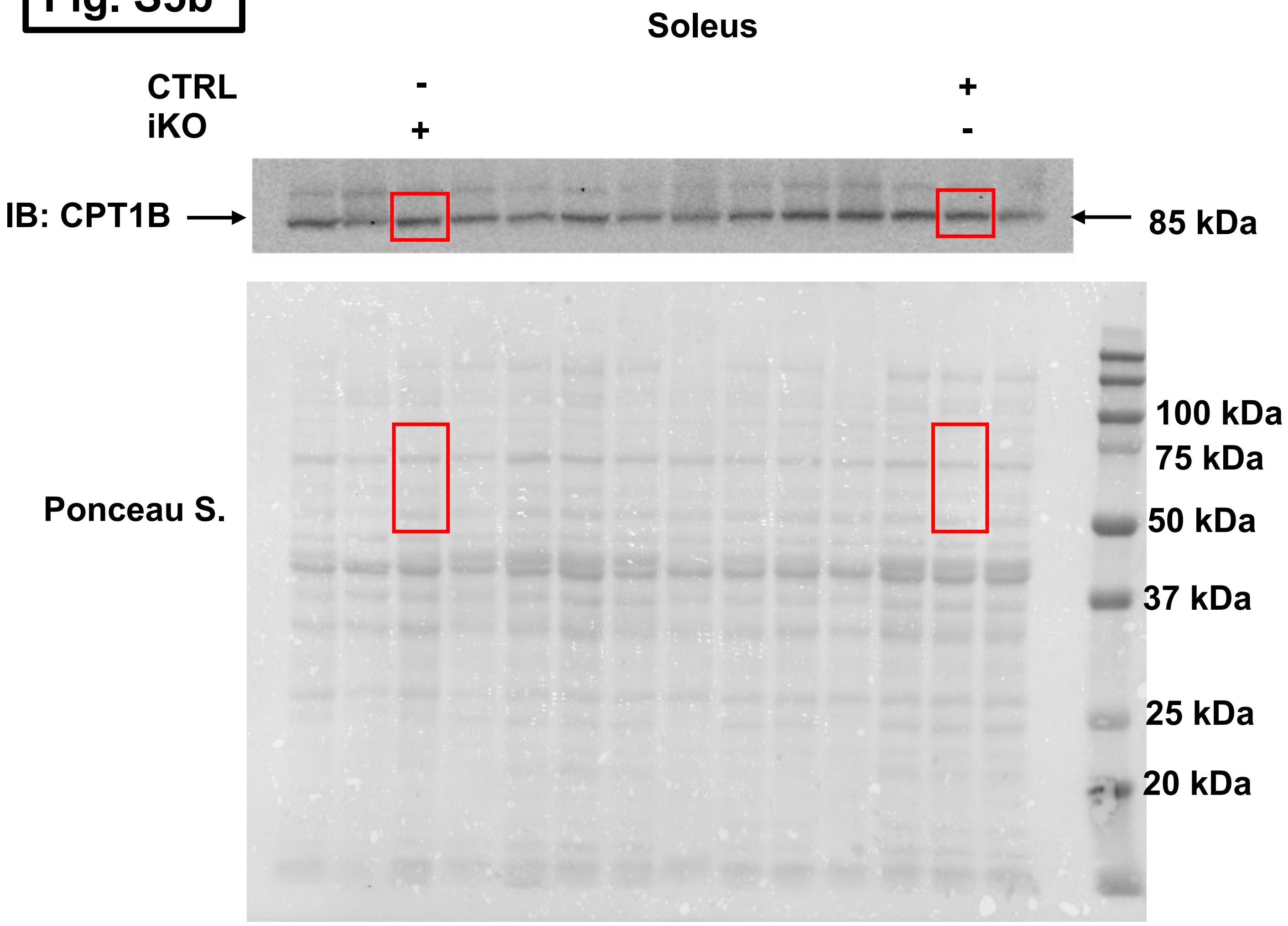

Soleus
CTRL - +
iKO + -
IB: CPT1B
85 kDa
Ponceau S.
100 kDa
75 kDa
50 kDa
37 kDa
25 kDa
20 kDa

Fig. S5b

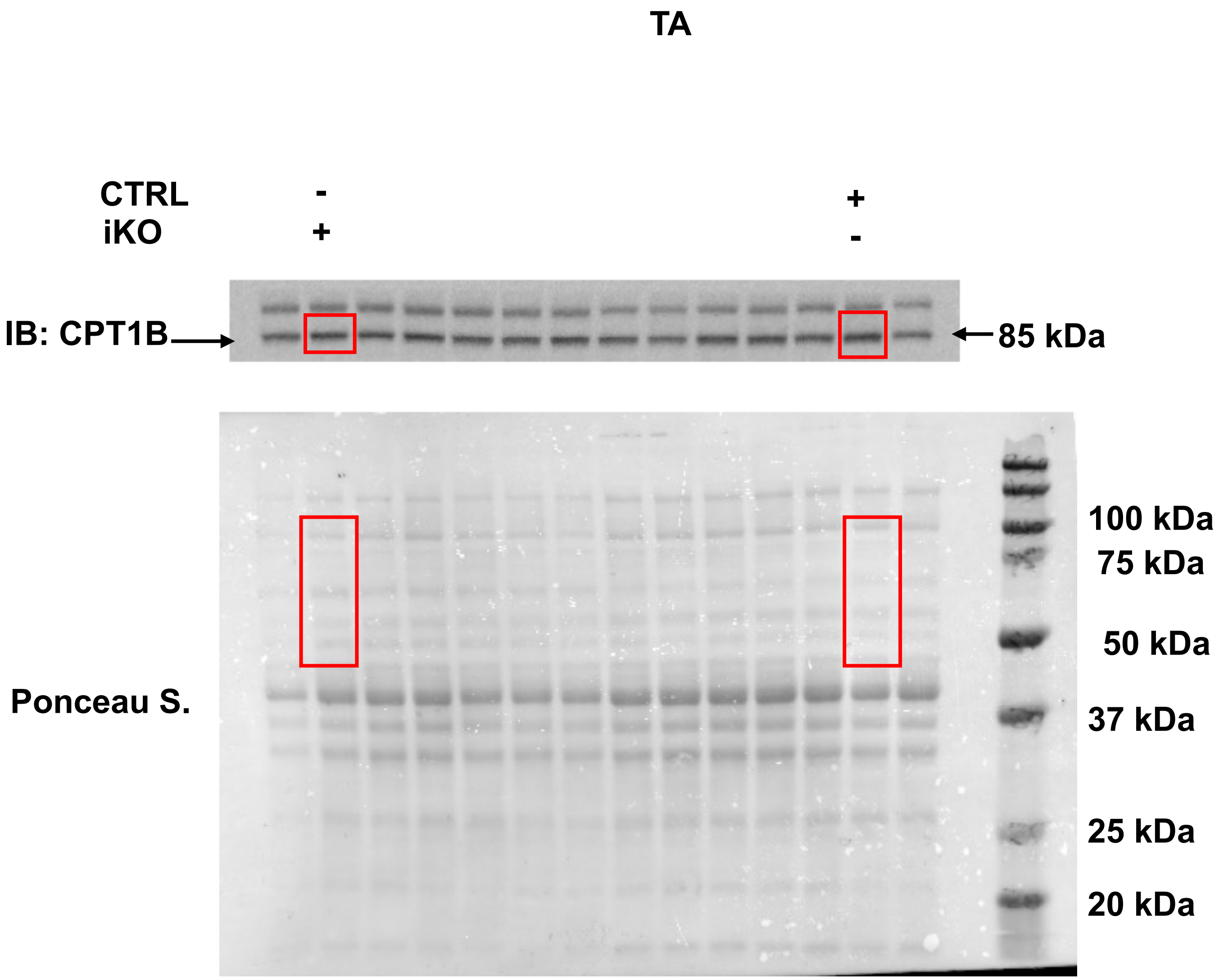
TA
CTRL
iKO
-
+
+
-
IB: CPT1B
85 kDa
Ponceau S.
100 kDa
75 kDa
50 kDa
37 kDa
25 kDa
20 kDa

**Fig. S5C**

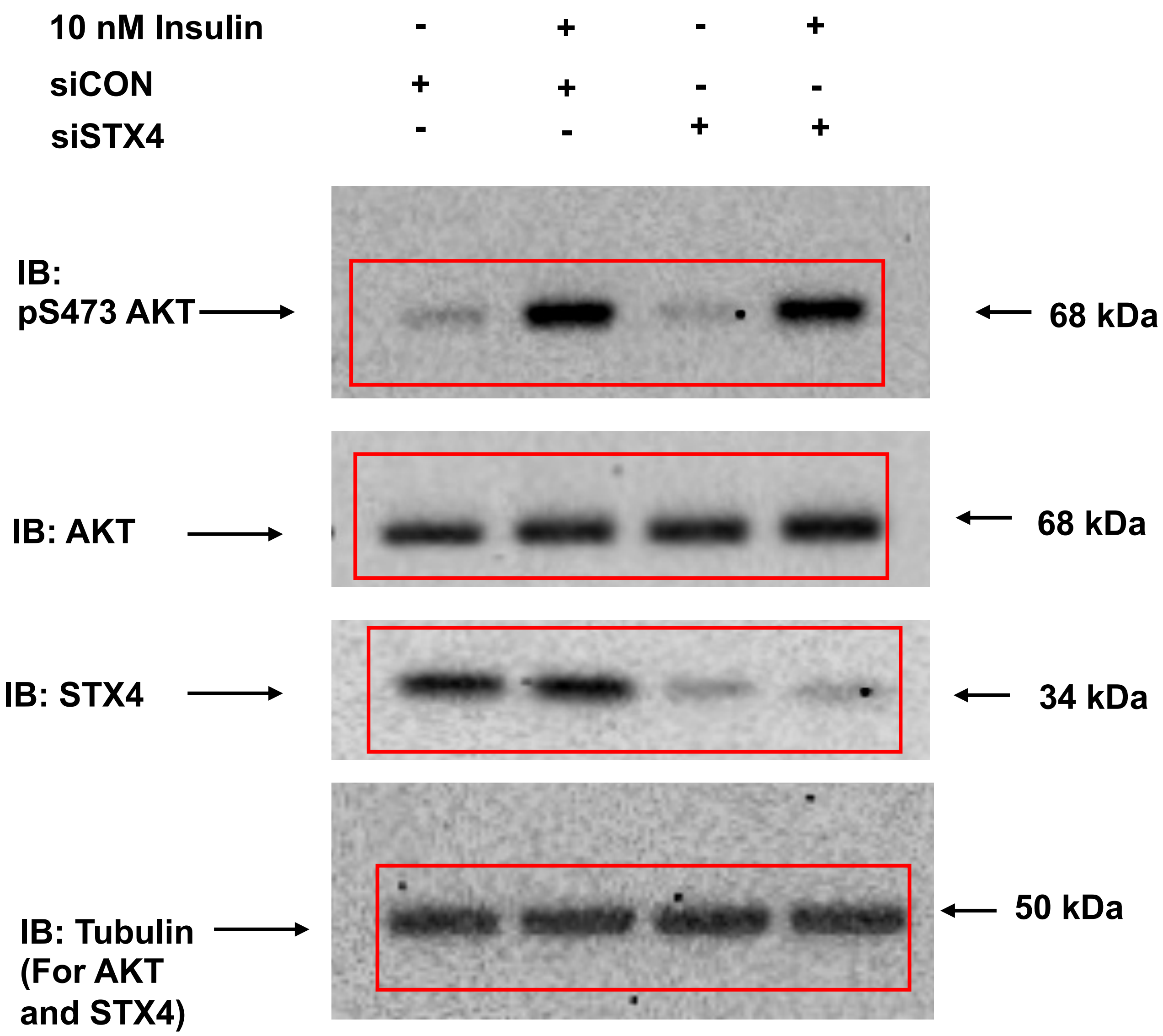
10 nM Insulin - + - +
siCON + + - -
siSTX4 - - + +
IB: pS473 AKT
68 kDa
IB: AKT
68 kDa
IB: STX4
34 kDa
IB: Tubulin (For AKT and STX4)
50 kDa
